# Fitness effects of antimicrobial resistance genes in changing environments

**DOI:** 10.64898/2026.03.06.710025

**Authors:** Alberto Hipólito, Lucía García-Pastor, Alexandra von Strempel, Anna S. Weiss, Jorge Sastre-Domínguez, Ester Vergara, Julio Álvarez, Ayari Fuentes-Hernández, Rafael Peña-Miller, Álvaro San Millán, Bärbel Stecher, José Antonio Escudero

## Abstract

The evolutionary success of antimicrobial resistance (AMR) genes is generally seen as a trade-off between their function in the presence and their cost in the absence of antibiotics. Mobile integrons are genetic elements that recruit and disseminate dozens of AMR genes among Gram-negative pathogens. Here, we have measured the fitness effects of 136 integron genes conferring resistance against several antibiotic families. We have found a significant proportion having neutral and positive effects in the absence of antibiotics. We confirmed this using a mouse model, where we also observed cases of changes in the sign of fitness effects. This led us to unveil that oxygen availability modulates the cost of AMR genes. Using a stochastic model, we show that fluctuating aerobic/anaerobic conditions can rescue AMR genes in the absence of selective pressure. Here we provide a comprehensive analysis of the cost of AMR at the gene level challenging the traditional fitness-resistance trade-off hypothesis.

## Introduction

Antimicrobial resistance (AMR) is a major challenge to health globally, causing between 1.2 and 5 million deaths per year^1,2^. The extensive use of antibiotics, together with the horizontal transfer of mobile genetic elements (MGE) such as plasmids, transposons and integrons, have fostered the rise of AMR^3,4^. However, the acquisition and expression of resistance determinants generally entails a fitness cost in the absence of selective pressure. This cost is a key player in the evolution of AMR, as it is considered the main driver of resistance reversibility at the community level^5^. The cost of AMR has often been studied for whole plasmids^6,7^, where resistance genes represent only a small fraction of the content but account for a significant part of the burden imposed to the host^8^. Nevertheless, conflicts between genomes and MGEs can also influence the cost of resistance^5,9,10^.

There is also a wealth of studies addressing the cost of AMR at the gene level, but the diversity of genetic context for these genes makes difficult to deliver studies that are broad, comparable and biologically sound at the same time. An interesting exception to this rule is the case of mobile integrons (MIs), genetic elements that capture and stockpile genes of adaptive value for bacteria. These genes are encoded in small mobilizable elements called integron cassettes (ICs). ICs are inserted by the integrase at the insertion site in the integron platform (*attI*), forming an array of variable functions^11–13^. ICs are generally promoterless and their expression is driven by the Pc promoter located within the platform, adjacent to the *attI*. When in first position of the array, the level of expression of ICs will depend exclusively on the Pc promoter^14^, while in downstream positions, expression levels are modulated by polar effects exerted by other cassettes upstream^15^. Carried on conjugative plasmids, MIs serve as vehicles for 177 antimicrobial resistance cassettes (ARCs) against 12 antibiotic families^16,17^, and are important drivers of multidrug resistance among Gram negative clinical pathogens^18–20^. Additionally, they also encode bacteriophage resistance cassettes, highlighting their role in adaptation^21^.

Genes in integrons share a common genetic background and set of regulatory rules^16,22^. Yet the abundance of each ARC varies significantly -as reflected in the INTEGRALL database^23^-even among those providing the same resistance levels. This suggests that evolutionary forces other than selective pressure on their function (antibiotic use) must be playing an important part in the success of ARCs. We hypothesize that the fitness cost must be one of these forces. While it has been shown that the cost of the integron platform is low^24^, little is known about the fitness cost of individual ARCs. Here, we take advantage of the pMBA collection, a set of isogenic *E. coli* strains containing 136 different cassettes in first position of a class 1 integron^16^ to characterize the cost of ARCs. Through competition experiments, we found an unexpectedly large proportion of genes that were neutral or beneficial in the absence of antibiotics. To understand the validity of our laboratory results in more realistic conditions, we performed competitions in a mouse model. Our data confirms that ARCs can be beneficial in the absence of selective pressure both *in vitro* and *in vivo*, acting as an exception to the proposed fitness-resistance trade-off. Additionally, we demonstrate that environmental conditions in the animal model can change the magnitude and the sign of the fitness effect of certain ARCs, highlighting their context dependence. We disentangled the differences between *in vivo* and *in vitro* environments to reveal that oxygen availability -a key changing condition in the lifestyle of enterobacteria- plays a key role in modulating fitness cost. To explore how environmental variation shapes the long-term dynamics of ARCs, we developed a stochastic competition model parameterized by our empirical data. This allowed us to simulate population trajectories under constant and fluctuating conditions, revealing that trade-offs across environments can lead to the evolutionary rescue of genes that would otherwise go extinct in a constant environment.

Altogether, this work provides a broad, comparable and biologically sound analysis of the cost of AMR. We reveal that the cost of genes is influenced by oxygen availability, a commonly changing condition for bacteria in the gut. Our results pave the way for a better understanding of the ecology and evolution of resistance.

## Results

### ARCs exhibit diverse fitness effects in *E. coli*

We sought to estimate the fitness effect of the 136 ARCs comprised in the pMBA collection in *E. coli* MG1655. The pMBA vector contains a class 1 integron mimicking the natural genetic environment of ARCs when in the first position of an array. Their expression is granted by a strong version of the Pc promoter (PcS) and they are followed by a *gfp* gene as a second cassette. The ARCs in the collection confer resistance against 8 families of antibiotics, including the most clinically relevant. This allows to study the fitness effect of each ARC in isogenic conditions and in their native genetic context (Figure 1A)^16^. When analyzing the growth curves of each pMBA strain, variations in growth rates become evident (Supplementary Figure S1). To better quantitate these differences, we used flow cytometry to conduct pairwise competitions *in vitro* in the absence of selective pressure. This technique is based on differentiating populations based on fluorescence, and is highly precise, detecting fitness differences as small as 1%^25,26^. To use it, we competed pMBA_ARC_ derivates (*gfp*^+^) with a strain carrying a *gfp^-^* empty vector (pMBA_□_ *gfp*^-^) (Figure 1B). Yet, comparisons are not straightforward in this setting, because recent studies from our lab have shown that cassettes modulate the expression of genes downstream^15^. Hence, the cost of the *gfp* might not be constant in our collection and this needs to be taken into account. To do so, we competed 17 pMBA_ARC_ strains showing different GFP levels, against their respective *gfp^-^*mutants. This led us to obtain the regression line that describes the relationship between expression and cost of the GFP (Figure 1C). This allowed us to subtract the cost of the GFP in all competitions and to obtain the cost of the 136 ARCs without confounding effects (Figure 1D and Supplementary Table 1). Supported by a linear regression model we determined that only 48% of ARCs (65/138) are detrimental for bacterial growth in the absence of antibiotics (with some being too costly to be quantitated using pairwise competitions). The rest of ARCs are divided equally between those that do not entail a significant cost and those that confer a significant fitness gain to the host in the absence of selective pressure (26% (36/136) each group) (Figure 1D). The most extreme case is that of *ereA2* which confers up to 39% of growth benefit. We could observe clear differences between the fitness effects of antimicrobial resistance families (Figure 1E and Supplementary Table 2). We focused on the three most abundant families: trimethoprim (*dfr*), aminoglycoside *(aa, aph, sat*), and ß-lactam (*bla*) resistance genes. While trimethoprim and aminoglycoside resistances genes follow a normal distribution centered at 1, ß-lactamases have a significant tendency towards high cost, in accordance with other reports^6,27^, with only some exceptions showing positive fitness effects.

**Figure 1.**
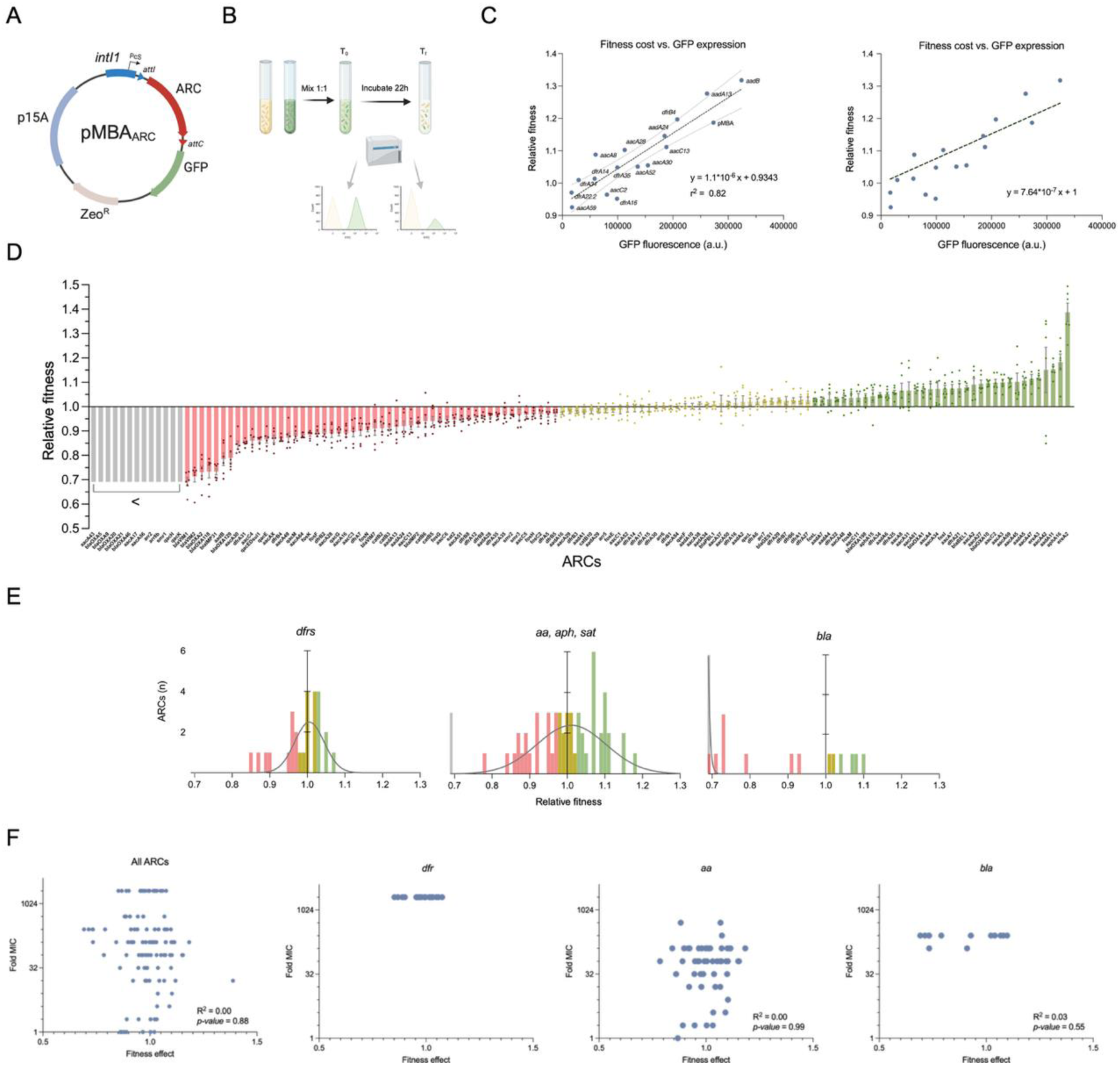
ARC fitness effect characterization. **A)** Schematic representation of pMBA_ARC_ vector showing the integrase with the PcS and the recombination site (*attI*), the ARC encoded is followed by a *gfp* gene, a zeocin resistance marker and the p15A origin of replication. **B)** Diagram of competition assays using flow cytometry (modified from ^26^). **C)** Correlation between fitness cost and GFP intensity of 17 different pMBA derivates. Simple linear regression is represented as a dotted line together with the 95% confidence interval (CI). r^2^ = 0.82. Each point represents the mean of three independent replicates. Right panel shows the linear regression forcing y = 1 the intercept. **D)** Graph showing the relative fitness of each ARC-carrying clone compared to the ARC-free isogenic clone in the absence of antibiotics (after correction for GFP expression). Bars represent the mean of 6 independent replicates, error bars represent the standard error of the mean (SEM). Red, yellow and green bars represent costly, neutral and beneficial fitness effects of ARCs as determined statistically. Grey bars represent ARCs too costly to be measured. **E)** Histograms showing data from D) grouped by antibiotic family: *dfrs* (trimethoprim), *aa, aph,sat* (aminoglycosides), and *bla* (ß-lactams). Grey lines represent gaussian fits to the data. **F)** Correlation between fitness effect and MIC increase of all ARCs, *dfrs*, *aa*, and *bla* families. Fold MIC data is collected from^17^. Pearson R^2^ and p-value are indicated in each graph. Correlation could not be calculated for *dfrs* for the lack of variation in the dependent variable.

A key assumption of the resistance/cost trade-off hypothesis is that the cost of genes is related to the level of resistance they confer. To test this, we used data form a previous work where we had characterized the levels and profiles of resistance for the whole pMBA collection against a variety of molecules from each antibiotic family^17^. For each gene, we correlated the maximum fold-change in resistance with its fitness effects. We found that correlations were virtually inexistant for all genes either when measured together or when grouped by families (R^2^ = 0.00 for all genes and for *aa*; incalculable for *dfrs* and R^2^=0.03 for *bla*) (Figure 1F). This data challenges the universality of the resistance/cost trade-off hypothesis and rather supports a resistance-dependent effect on fitness.

### ARCs can provide fitness gains *in vivo*

Compelled by the abundance of positive fitness effects in our collection, and the extreme cases like *ereA2*, we sought to confirm these results *in vivo.* To do so, we used a controlled-microbiota mouse model in which mice are stably colonized by the OMM^12^ bacterial consortium, composed of twelve species of the five major eubacterial phyla of the murine gastrointestinal tract^28^. This model is widely used to study microbial interactions and enteric infections^29,30^ (Figure 2A). We selected three of the most beneficial ARCs *in vitro* (*ereA2*, *aacA7*, and *bla*_OXA-10_) (depicted in Figure 3A) to perform competition assays *in vivo* (in the gut of mice) during 7 days in the absence of antibiotics (Figure 2B). In this experimental setting, populations are not distinguished on the basis of fluorescence, so we knocked out the *gfp* in both strains to avoid its confounding effect on cost (pMBA_ARC_ *gfp*^-^ *vs*. pMBA_□_ *gfp*^-^). We inoculated a 1:1 mixed culture at day 0 and collected fecal samples daily until day 4 and at day 7, when mice were sacrificed and cecal contents were collected. We determined total *E. coli*/pMBA load and relative proportions of competing strains along the experiment (Figures 2C and 2D). As a control, we examined the composition of the OMM^12^ consortium in fecal and cecal samples using qPCR and did not observe differences between days 0 and 7 in the different mice groups (Supplementary Figure S2), suggesting that the competition did not disturb the microbiota.

**Figure 2.**
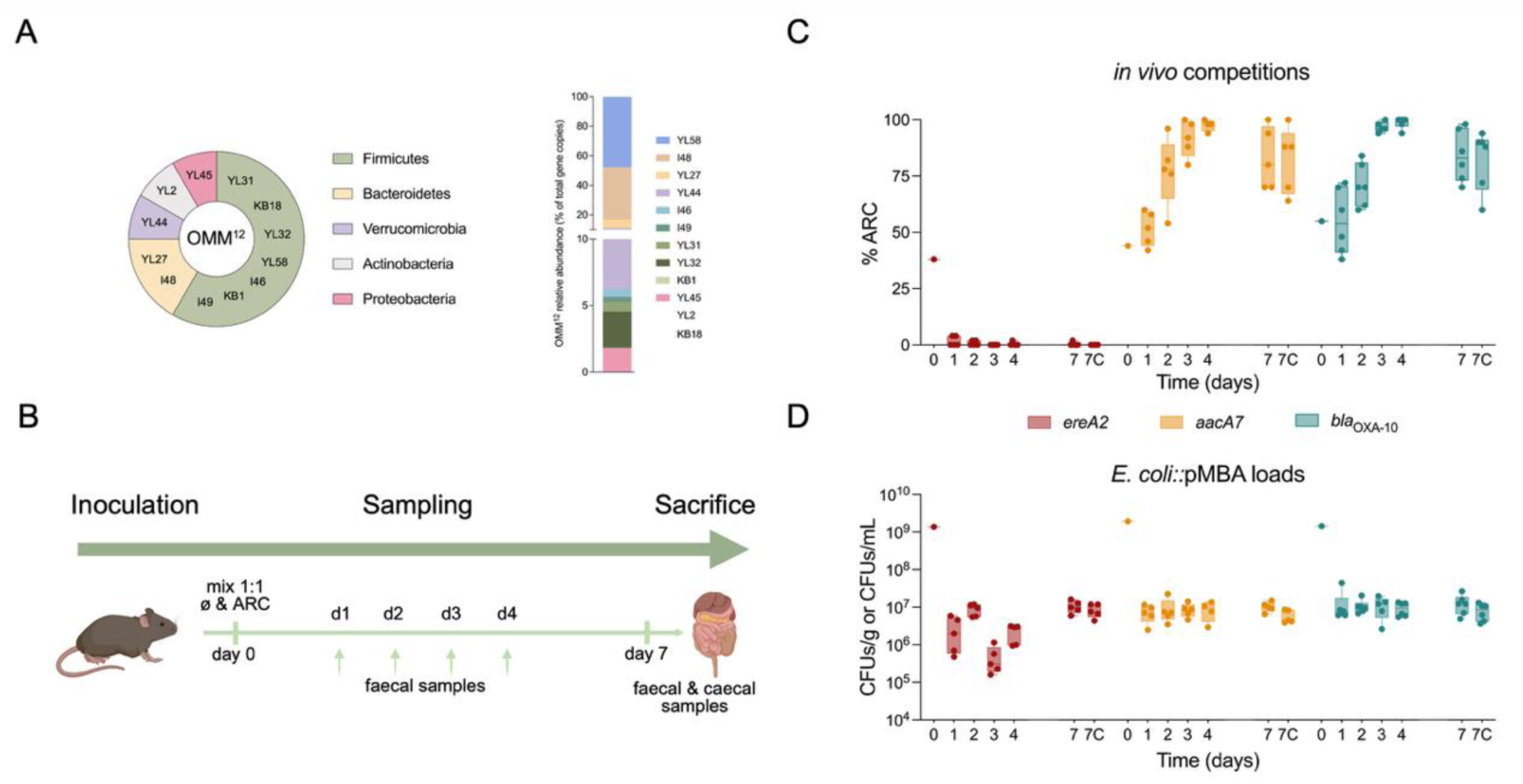
*In vivo* quantitation of ARCs fitness effect. **A)** Experimental design. OMM^12^ gnotobiotic mice were inoculated with a 1:1 mix (pMBA_ARC_ ΔGFP : pMBA□ ΔGFP). Fecal samples were taken daily from day 0 to 4. At day 7 mice were sacrificed and fecal and cecal samples were also taken. **B)** Oligo mouse microbiota OMM^12^ composition and relative abundance in a representative mouse at the beginning of the experiment (YL2 and KB18 are not present in the graph as their levels are under the limit of quantification). **C)** pMBA_ARC_ relative abundance compared to pMBA□ in the bacterial mixtures *in vivo* during 7 days in feces and cecal content (7C). **D)** *E. coli*/pMBA loads in feces and caecum quantitated as CFUs/g or CFUs/mL. Each competition is represented with a different color: red (*ereA2*), yellow (*aacA7*) and blue (*bla*_OXA-10_). Data is represented using a box-and-whisker plot, where each experimental group consists of at least five individuals represented by dots. Horizontal lines inside boxes indicate median values, upper and lower hinges correspond to the 25^th^ and 75^th^ percentiles, and whiskers extend to the minimum and maximum values

**Figure 3.**
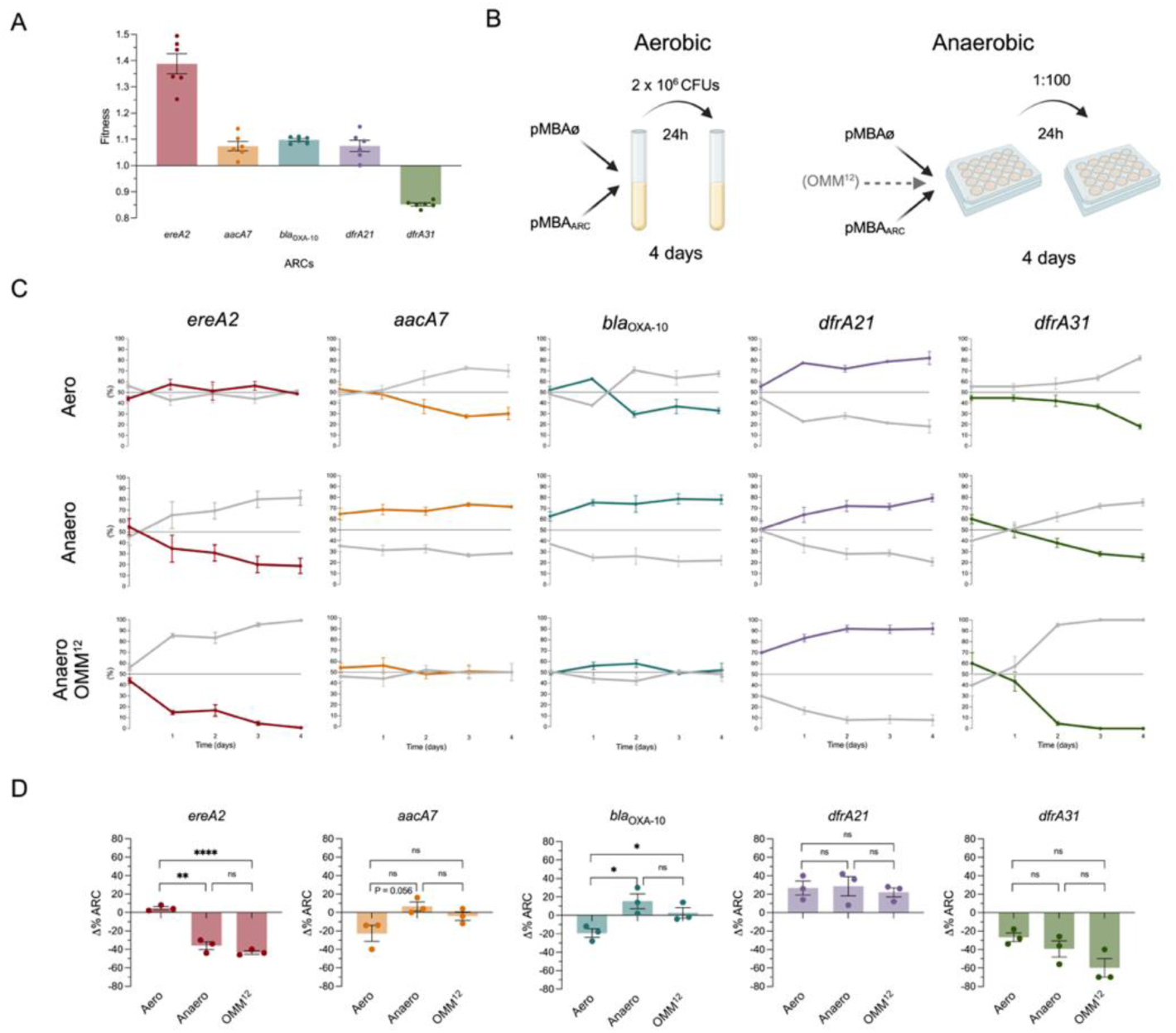
Long-term *in vitro* competition assays under different environmental conditions. **A)** Fitness effect measured by flow cytometry competition assays (data from Figure 1) of a subset of ARC: *ereA2* (red), *aacA7* (orange), *bla*_OXA-10_ (blue), *dfrA21* (purple) and *dfrA31* (green). **B)** Experimental design of both aerobic and anaerobic long-term competition assays. **C)** Quantitation of ARC population in the competitions during 4 days in aerobic (top panel), anerobic (middle panel), and in the presence of OMM^12^ consortium (anaerobic conditions and enriched media) (bottom panel). Each time point represents the mean of three independent replicates, error bars represent the standard error of the mean (SEM). Colored lines represent the pMBA_ARC_ ΔGFP populations while grey lines stand for pMBA□ ΔGFP. **D)** Variation in the population harboring ARCs (Δ%ARC) at the end of the competitions. Statistical differences were assessed using a two-sided unpaired t-test with Welch’s correction. * P < 0.05, ** P < 0.005, **** P < 0.0001. ns = non significative.

Our results showed a clear fitness advantage for strains carrying *aacA7* and *bla*_OXA-10_ in the absence of antibiotics. The *E. coli/*pMBA load (the sum of both competing strains) remained stable throughout the experiment at around 10^7^ CFU/mL or CFU/g yet the relative proportion of competitors changed. By day 4, in both cases the population containing the ARC had almost replaced the one with the empty vector, showing >90% prevalence. At day 7, the competitive advantage over the control was maintained, although less markedly, reaching 80% for *aacA7* and 70-80% for *bla*_OXA-10_ (Figure 2C yellow and blue boxes).

In contrast, the strain carrying *ereA2,* that provided the highest fitness gain *in vitro*, was quickly outcompeted after 1 day *in vivo,* suggesting a large cost (Figure 2C red boxes). Total *E. coli*/pMBA counts were lower and somewhat variable during the first days but normal by day 7 (Figure 2D). Variability in counts was not a consequence of a colonization defect, but rather due to plasmid loss in the *ereA2* population, which can be accelerated by costly genes (Supplementary Figure S3).

Our results suggest that certain ARCs are under positive selection in the gut, in the absence of antibiotic pressure. They also highlight that, at least in some cases, *in vitro* head-to-head competitions might not reflect precisely the cost of AMR *in vivo*.

### Anaerobiosis modulates the fitness effect of ARCs

Results from competitions *in vivo* with *ereA2* suggest that environmental conditions can change the sign of fitness effects of ARCs. We found this observation compelling and sought to determine the underlying cause of such change. Among the many differences between our *in vivo* and *in vitro* experiments, certain aspects cannot be tested *in vitro*, like the presence of immune components, enzymes and complex spatial structure. Yet others, like the lower oxygen concentration of the gut, or the presence of a microbial community can easily be tested *in vitro*. To do so, we selected the three ARCs tested in mice and added 2 more (*dfrA21* and *dfrA31*), to cover a variety of fitness effects in flow cytometry competitions (Figure 3A). Next, we performed long-term competitions (see materials and methods) in aerobic and anaerobic conditions, and in the presence or the absence of the controlled OMM^12^ microbiota (Figure 3B).

In aerobic conditions and in the absence of microbiota, the fitness effects of *ereA2*, *dfrA21* and *dfrA31* followed a similar trend as in flow cytometry (both experiments were done in MH, see Materials and Methods). Nevertheless, *aacA7* and *bla*_OXA-10_ were somewhat more costly at initial timepoints, although this cost leveled off by days 3 and 2 respectively. In anaerobic conditions in MH, *ereA2* became highly costly, while *aacA7*, and *bla*_OXA-10_ had more positive fitness effects, in accordance with the results obtained *in vivo*. On their side, *dfrA21* and *dfrA31* seemed unaffected by anaerobiosis, following similar dynamics in both environments. We also performed competitions in the presence of the OMM^12^ consortium, which needs anaerobic conditions and requires an enriched media, and did not observe an appreciable impact of the consortium compared to anaerobiosis only (Figure 3C). To obtain a clearer picture, we compared the change in relative abundance of ARCs between initial and final timepoints (Δ%ARC) (Figure 3D). We observed significant differences between aerobic and anaerobic conditions for *ereA2* and *bla*_OXA-10_, as well as a clear trend for *aacA7* (p-value = 0.056). All three genes changed the sign of the fitness effect between conditions. Instead, *dfrA21* and *dfrA31* showed consistent fitness effects in both conditions. This analysis confirmed that the OMM^12^ microbiota had no effect on our competitions. As a control, we verified potential fluctuations in the OMM^12^ consortium composition throughout the experiment and found no relevant changes generally, except for a mild decrease in *Clostridium innocuum* (I46) abundance in *dfrA21* and *dfrA31* competitions (Supplementary Figure S4).

Altogether, these data suggest that oxygen availability is an important factor influencing fitness effects of certain AMR genes. To confirm this, we conducted the whole set of *in vitro* flow cytometry competition assays but this time under anaerobic conditions (Figure 4 and Supplementary Figure S5 and Supplementary Table 3). This yielded a completely different landscape of fitness effects where only 4% of ARCs confer a fitness advantage without selective pressure (5/136), compared to 26% under aerobic conditions. Specifically, *arr7* is now the ARC providing the highest fitness gain (only 4.5% vs. 39% of *ereA2* in aerobic conditions) (Figure 4A and B green bars). In addition, the proportion of ARCs that do not entail a significant cost increased to a 35% (47/136) (Figure 4A and B yellow bars) while costly genes were now the largest group (62%, 84/136) (Figure 4A and B red and grey bars). At the family level, we observed a narrower distribution of fitness effects and a shift towards cost was evident in the trimethoprim and aminoglycoside but not in the ß-lactam resistance family (Figure 4C and Supplementary Table 4). Again, the fitness cost of ARCs in anaerobiosis did not correlate with the level of resistance conferred by the gene in this condition (data from^31^) (Figure 4D).

**Figure 4.**
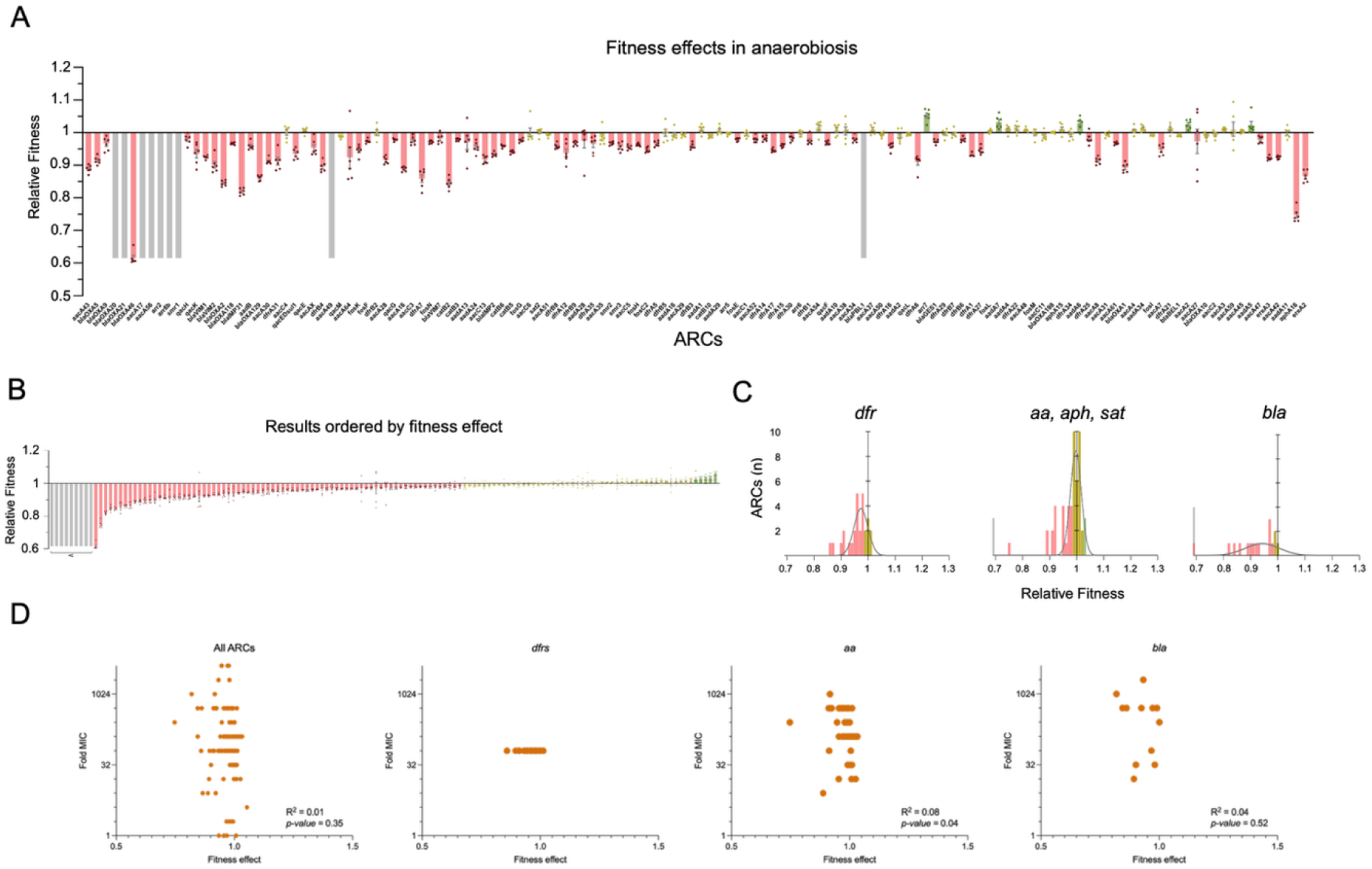
ARC fitness effect characterization in the absence of oxygen. **A)**. Graph showing the fitness effect of each ARC on bacterial growth in anaerobic conditions ordered as in Figure 1D. Bars represent the mean of 6 independent replicates; error bars represent the standard error of the mean (SEM). **B)** Graph showing data from panel A ordered from costly (left) to beneficial (right). **C)** Histograms show fitness distribution of ARCs from *dfr*, *aa*, and *bla* families. Grey line represents a gaussian fit to the data. Grey bars represent ARCs unable to be measured, red bars costly ARCs, yellow bars represent ARCs with no significant cost (according to a linear regression model), and green bars represent beneficial ARCs without selective pressure. **D)** Correlation between fitness effect and MIC increase in anaerobiosis of all ARCs, *dfrs*, *aa*, and *bla* families. Fold MIC data is collected from^31^. Pearson R^2^ and p-values are indicated in each graph.

The changes in the distribution of fitness effects led us to perform a more detailed analysis to understand if protein families are affected differently by anaerobiosis. When considering all ARCs together, we confirmed the narrower distribution of effects and their shift towards cost (Figure 5A). This change was driven by trimethoprim and aminoglycoside resistance genes (Figure 5B) but not ß-lactam and fosfomycin resistance genes, that were not affected (Figure 5C) (Supplementary Table 5). Nevertheless, a gene-by-gene analysis showed that anaerobiosis affected the fitness effect of 53% of ARCs (64/120) (Supplementary Table 6). Certain genes exhibit drastic changes in fitness effects that include changes in sign, like *ereA2* that goes from highly beneficial in aero- to costly in anaerobiosis; or *qacE,* that follows the opposite trend (Figure 5A red and yellow dotted lines). Despite the generalities observed for gene families, every family has at least one member significantly influenced by oxygen availability and most have a member changing sign. Even in the *bla* and *fos* families - which, at the family level, appear unaffected - most individual ARCs still show significant fitness changes depending on the presence or absence of oxygen (Figure 5C) (Supplementary Table 6). Hence, we conclude that oxygen availability impacts the fitness effects of AMR in a gene specific manner, but with recognizable effects at the family level. Oxygen pressure is a constantly changing condition for enterobacteria that thrive in the gut and are shed to the environment. Our results suggest that, on the long-term, the evolution of AMR might depend to some extent on fluctuating oxygen conditions.

**Figure 5.**
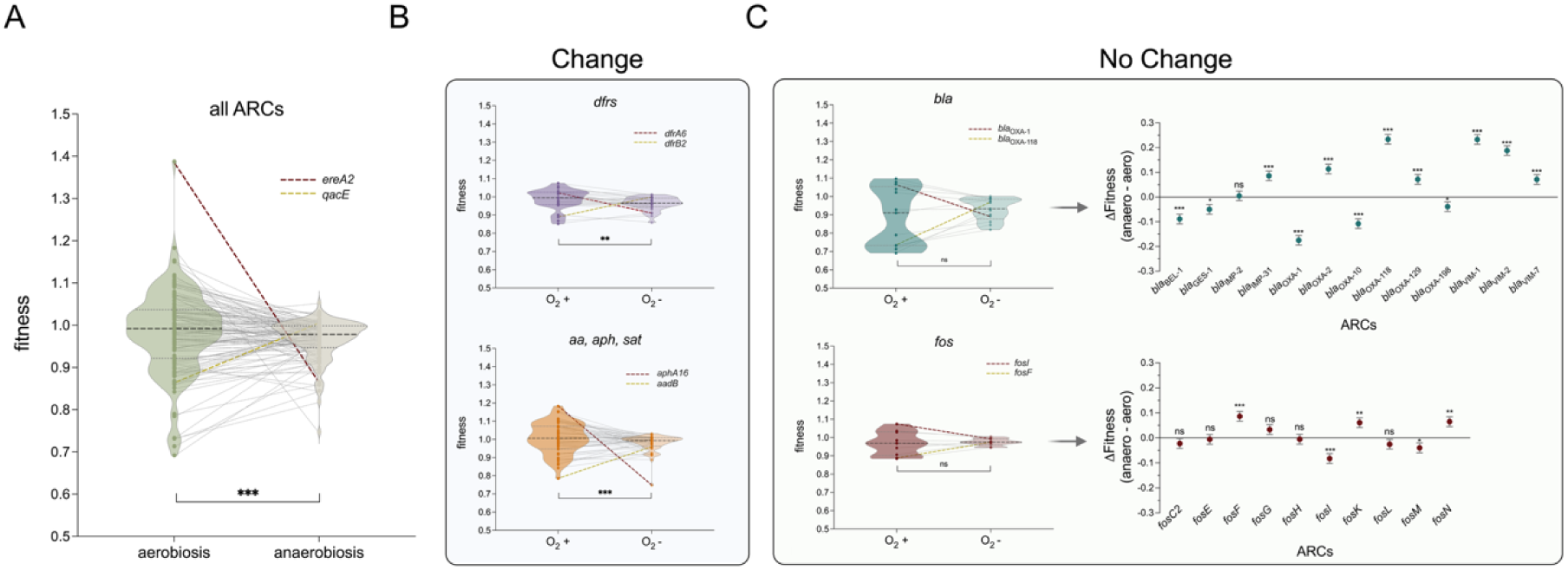
Comparison of fitness effects of ARCs under aerobic and anaerobic conditions. Violin plots show fitness effects in aerobiosis (dark color) and anaerobiosis (light color) of **A)** all ARCs, **B)** *dfrs* and aminoglycoside resistance genes, **C)** *blas* and *fos* (Data from Figures 1 and 4). Gray lines link the dots representing the same gene in both conditions. Red-dotted line represents relevant ARCs that are costlier in anaerobiosis while yellow-dotted line stands for ARCs lowering their fitness burden in the absence of oxygen. Panel **C** also show the difference in fitness effects of *bla* and *fos* members respectively. Statistical significance was assessed using a linear regression model. *: P<0.05; **: P<0.005; ***: P<0.0005. ns = non-significant.

### Environmental Rescue of ARCs During Oxygen Transitions

To further probe the role of oxygen fluctuations in ARC maintenance in the long term, we developed a stochastic competition model to simulate long-term pairwise competitions between ARC-bearing strains and the isogenic plasmid-empty control. Each daily growth cycle was defined as either aerobic or anaerobic using growth parameters calibrated from *in vitro* assays (see Methods and Supplementary Information). We obtained paired estimates of relative fitness from experimental competitions, defining a joint empirical distribution for each resistance family. We modeled this distribution using a bivariate normal, capturing the observed means, variances, and correlation between conditions. Synthetic ARC populations were then generated by sampling from these family-specific distributions and calibrating model parameters to match the sampled fitness values. This approach reproduced both the diversity and structure of the empirical fitness landscape, including variation in environmental responses across ARC families (Figure 6A).

**Figure 6.**
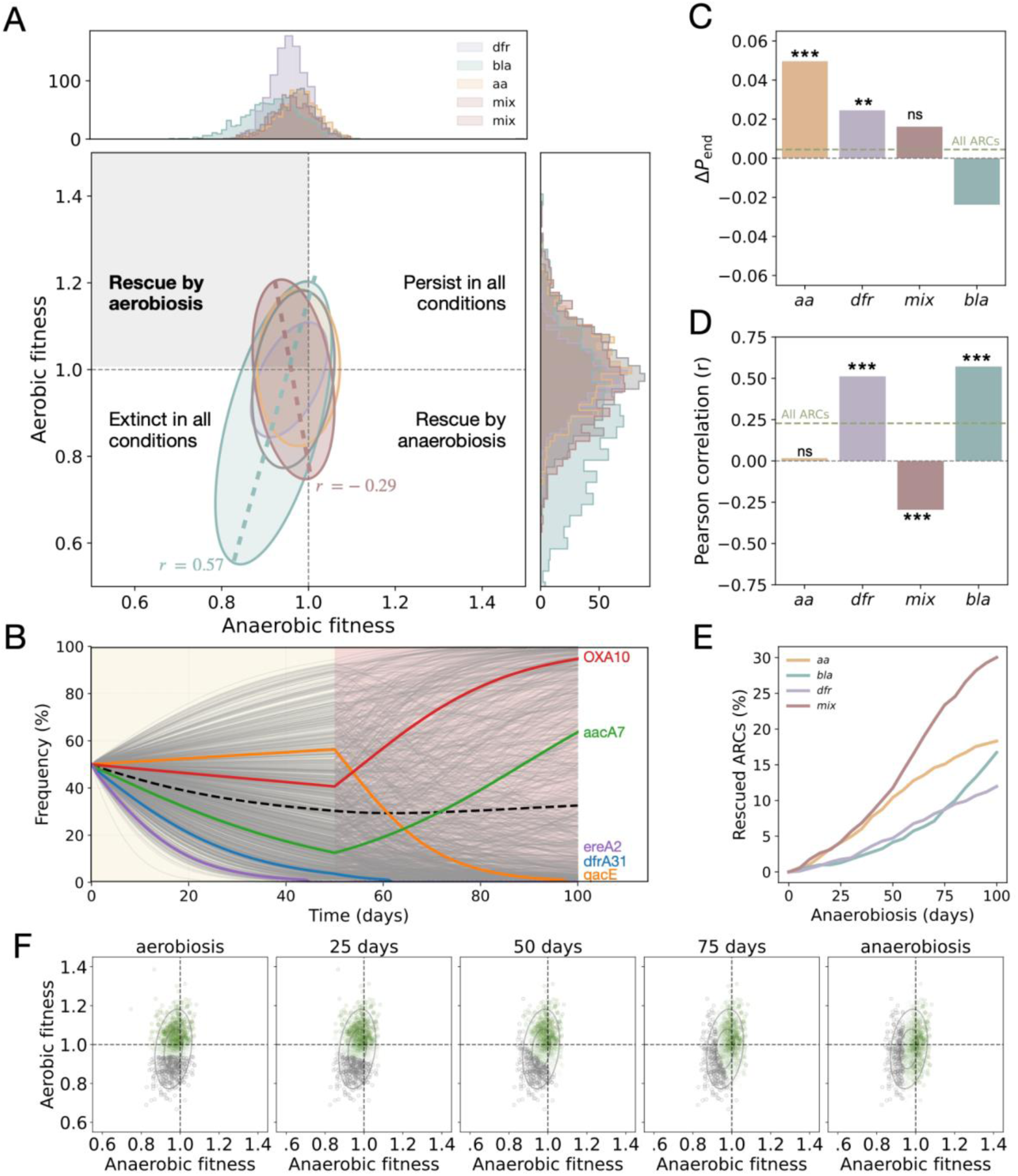
Modeling the evolutionary rescue of ARCs under fluctuating oxygen. **A)** Covariance ellipses (2 standard deviations) represent each ARC family’s fitness distribution, derived from the synthetic library used in the simulation model. Colors denote ARC families: trimethoprim- (dfr, purple), ß-lactam- (bla, green), aminoglycoside- resistance (aa, orange), and a group of unrelated genes against several families of antibiotics (mix, red). Marginal histograms display anaerobic fitness (top) and aerobic fitness (right) distributions. Dotted dashed lines indicate neutrality (fitness = 1). Strains in the top-right quadrant are fit in both environments and expected to persist; those in the bottom-left are unfit and prone to extinction. Off-diagonal regions contain strains with condition-specific advantages, enabling potential rescue upon oxygen shifts. **B)** Simulated frequency trajectories of ARC-bearing strains under a single anaerobic-to-aerobic switch at day 50. Each line shows one synthetic genotype; highlighted lines represent illustrative ARCs. The black dashed line shows the population average, which remains stable despite divergent individual dynamics. Some strains recover from early decline, while others drop despite early success, reflecting contrasting environment-specific fitness profiles. **C)** Difference in final ARC survival under a single anaerobic-to-aerobic switch, relative to the average survival in constant conditions. Bars show ΔP_end_; positive values indicate increased persistence via environmental switching. Asterisks denote statistical significance (*: P<0.05; **: P<0.005; ***: P<0.0005. ns = non-significant). **D)** Pearson correlation between aerobic and anaerobic fitness within each ARC family. Stronger correlations indicate constrained trade-offs across environments; weaker or negative values reflect greater potential for environment-specific responses. **E)** Fraction of ARC strains rescued after an anaerobic-to-aerobic switch, as a function of anaerobic duration. Families with weaker or negative fitness correlations show higher rescue rates, consistent with broader trade-offs and increased sensitivity to environmental change. **F)** Fitness-space geometry of ARC persistence under different switch times. Each panel shows the fitness values of the synthetic ARC library in anaerobic (x-axis) and aerobic (y-axis) relative fitness space. Panels are ordered from left to right by the timing of a single switch from anaerobiosis to aerobiosis: constant aerobiosis, switch after 25, 50, and 75 days, and constant anaerobiosis. Green points indicate ARC-bearing strains that persist at the end of the simulation for the corresponding switch time, with transparency reflecting the frequency of persistence across replicate simulations. Rotated ellipses denote the covariance structure of fitness variation (1 and 2 standard deviations). Across panels, the orientation of the persistence boundary rotates with switch timing, reflecting the relative contribution of anaerobic and aerobic fitness to long-term survival. It is of note that the boundary does not intersect at 1 because the simulation stops at 100 days. This allows strains with very mild negative values to be retrieved.

This *in silico* framework allowed us to simulate serial transfer experiments under defined oxygen regimens, using the same daily growth and dilution structure as our laboratory passaging protocol (Figure 3B). We first examined constant environments as a baseline. Under continuous aerobiosis, ARC-bearing populations often persisted through the duration of the experiment (∼59<% survival), with ∼19% reaching fixation. In contrast, under continuous anaerobiosis, ∼43% of ARC populations went extinct, and fewer than 1% reached fixation, indicating that even surviving strains generally remained at low frequency. These results establish a baseline for ARC persistence under static conditions, where long-term survival depends entirely on fitness within a single environment. However, the life cycle of gut microbes often involve drastic fluctuations in oxygen availability, from the anaerobic gut to the aerobic environment. To explore how such dynamics influence ARC outcomes, we next simulated a single switch from anaerobiosis to aerobiosis and evaluated its impact on ARC dynamics.

In our 100-days simulations, we defined a ‘rescued’ ARC as one that would go extinct under constant anaerobic conditions but survives when an aerobic phase is introduced. Environmental shifts had minimal effect on the mean ARC frequency across simulations, but revealed striking heterogeneity in individual outcomes (Figure 6B). Some ARC-bearing lineages increased in frequency post-switch, while others declined, reflecting variation in aerobic–anaerobic fitness trade-offs of individual ARCs. For instance, *ereA2*, which incurred a large fitness cost under anaerobic conditions but provided a benefit under aerobic conditions, went extinct under continuous anaerobiosis yet was rescued when an aerobic phase was introduced before day 45 (Supplementary Figure S6). The proportion of rescued ARCs increased with the length of anaerobic exposure (Figure 6C; rescue rate after 20 days: ∼3%; after 80 days: ∼17%). When anaerobic phases were short, survival rates remained high (>50%), limiting the number of strains eligible for rescue. In contrast, longer anaerobic periods increased extinction risk, resulting in more frequent rescue events upon switching. This example underscores that an ARC must be sufficiently fit in at least one environment to persist: strains with relative fitness < 1 in both conditions invariably went extinct regardless of switching.

We compared endpoint survival after a single anaerobic-to-aerobic switch at the experiment’s midpoint (day 50) to the average survival in constant conditions; the difference (ΔP_end_) quantifies the effect of switching on ARC persistence. Families with weaker correlations between aerobic and anaerobic fitness, such as *aa* and *dfr*, showed the greatest gains in survival following the switch (ΔP_end_ = 0.05 and 0.025, respectively; Figure 6C). In contrast, *bla*, which exhibited a strong positive correlation, showed a decrease in survival (ΔP_end_ = – 0.024). Genes outside these families, comprising a heterogeneous set of resistance mechanisms (mix group, defined in Supplementary Table 7), showed the most pronounced fitness trade-off (r = –0.29, p < 0.001; Figure 6D) and the highest frequency of rescued strains (Figure 6E). To visualize how fitness distributions shape survival, we mapped ARC persistence onto the two-dimensional fitness space. In constant aerobiosis, the survival boundary is nearly vertical, while in anaerobiosis it is horizontal; intermediate switch times rotate this boundary, reflecting combined selection (Figure 6F; Supplementary Figure S7). Strains with fitness <1 in both conditions always went extinct. This geometric perspective highlights how the timing of environmental change interacts with the shape and orientation of fitness variation to determine ARC survival, consistent with our *in vitro* and *in vivo* findings that oxygen dynamics drives long-term resistance persistence.

## Discussion

Resistance determinants generally impose a cost to the host. This fitness-resistance trade-off drives the ecology and evolution of resistance, and shapes our measures to counteract it. Indeed, one of the most effective policies to combat AMR is the responsible use of antibiotics, that aims precisely at exploiting this trade-off. The cost of AMR has been demonstrated experimentally and varies in relation to the coding platform, bacterial host, genetics and environmental conditions^32–36^. The cost imposed by some plasmids comes mainly from the expression of antimicrobial resistance genes^5,8^. However, broad studies quantitating the fitness effect at the gene level are not common. In an ambitious work, Porse *et al*. studied in *E. coli* the fitness cost of 126 genes from Gram positive and negative bacteria^37^. To achieve this, they used a standardized cloning vector where genes were genetically decontextualized. Mobile integrons are an important source of antimicrobial resistance cassettes among the most relevant Gram-negative pathogens. Because integrons provide a common genetic background and set of regulatory rules for ARCs, they represent a great model to perform large and comprehensive studies on the fitness dynamics of AMR determinants. Here, we used an isogenic collection of 136 ARCs to study their fitness effects in different environmental conditions. Compared to Porse’s results, our data show an unexpectedly high proportion of ARCs that are beneficial to their hosts in the absence of selective pressure. This is an important addition to existing evidence challenging fitness-resistance trade-offs^38^. A key advantage of our collection is that it allows a transversal analysis. For instance, we can consider the highly homogeneous group of 22 *dfr* genes in our collection. These genes confer resistance to the same antibiotic (trimethoprim) to the exact same level^17^ and through the same biochemical mechanism (they are all dihydrofolate reductases), yet they entail a variety of fitness effects ranging from positive to negative (in aerobiosis). This not only challenges the assumption that AMR is inherently costly, but more importantly, it suggests that the cost may not be a consequence of the function of the gene (Figures 1, 2 and 4 and Supplementary Tables 1 and 3). It is possible that the cost derives from genetic conflicts between ARCs and the host’s genome, or to the presence of moonlighting activities of AMR genes. Previous works suggest that AMR genes that do not confer relevant resistance levels in their native host might be playing a role in bacterial signaling and metabolism, including the use of antibiotics as a carbon source^39–42^. It is therefore possible that, in our collection, ARCs conferring a fitness advantage are playing a secondary role in the absence of the drug, that contributes to essential processes of the host.

Independently of the underlying mechanism, our results from *in vivo* experiments confirmed that some genes -*bla*_OXA-10_ and *aacA7*- are indeed beneficial for bacterial growth in clinically-relevant environments in the absence of antibiotics. This is not only relevant for their role as AMR genes, but also because integrons can contain many genes and are found in a broad variety of plasmids carrying other AMR genes encoded elsewhere. Genetic linkage sets the stage for co-selection phenomena, so that ARCs with positive fitness effects could diminish plasmid cost indirectly stabilizing other AMR genes in populations (and viceversa). Studying the importance of this effect is crucial for understanding the spread of antimicrobial resistance and to tackle this global health challenge.

Based on discrepancies between *in vitro* and *in vivo* results, we have unveiled the influence of O_2_ in modulating fitness effects. In general, ARCs tend to be costlier in anaerobic conditions with a narrower distribution of fitness effects. However, a gene-by-gene analysis is needed due to the variety of scenarios (Figure 5 and Supplementary Table 6). We had previously shown the impact of O_2_ on the performance of AMR genes^31^ suggesting that this is a key environmental condition to consider in the field of AMR. This is especially the case for enterobacteria, whose lifestyle includes a constant switch between aerobic and anaerobic conditions. Additionally, oxygen is present in the gut under certain circumstances, like in the gut of newborns or possibly during inflammatory processes^43,44^. It is noteworthy that integrons carrying AMR genes are prevalent in 1 month old babies^45^. Our computational model shows that fluctuating oxygen availability reshapes the effective fitness landscape of ARC-bearing strains. In our simulations, environmental switching shifts the boundary that separates persistent from declining strains in the aerobic–anaerobic fitness plane, revealing how the timing of environmental change determines which fitness combinations support long-term persistence. Strains with similar mean fitness can nonetheless follow different trajectories depending on how their fitness values are distributed and correlated across environments. These results indicate that stability under switching regimes depends not only on absolute fitness but also on cross-environment covariance, providing a mechanistic explanation for why environmental transitions can maintain ARC diversity that would otherwise be lost under constant conditions.

Moreover, the differences in fitness effects between environmental conditions could be key in the ecology of these ARCs. Environments in which AMR genes have positive fitness effects might act as a reservoir of these genes for other environments, in a source-sink like dynamic. A good example of this is *ereA2*, the gene showing the most drastic changes in fitness. *ereA2* encodes an erythromycin esterase and its presence in integrons is puzzling because Gram-negatives are naturally resistant to erythromycin, making implausible its selection through antibiotic use. The large fitness gain it confers in aerobiosis might serve as an explanation to its presence, conferring an evolutionary advantage in aerobic environments. Nevertheless, its high cost in anaerobic environments and mice models should represent a burden to its spread in clinical settings. This could explain why *ereA2* is not extremely prevalent, and is often found truncated in databases, where clinical isolates – often thriving in anaerobic environments - are overrepresented compared to environmental samples. We applied a similar hypothesis to account for the differential prevalence of individual ARCs, expecting that ARCs with higher costs would be rarer in databases, while beneficial ARCs would be more common. A good example of such correlation is *aadA5*, the most reported ARC in class 1 MIs in *E. coli* that shows beneficial effects in all our experiments (11% and 2% in aerobiosis and anaerobiosis, respectively). However, such clear correlations were not observed with other cassettes in the collection. This can be due peculiarities of MIs, where polar effects of cassettes might play a role in modulating the fitness effects of other^15,21^. Additionally, other environmental factors, including genetic background, spatial structure or the varying strengths of antibiotic selection, are likely at play in real conditions (i.e.: in the databases). They change fitness landscapes in a myriad ways generating a *seascape* of yet unknown navigability^46–48^. Hence, finding a direct correlation between fitness effects of ARCs in the lab and their prevalence in databases is not yet feasible. Expanding this study to a broader range of clinically relevant species and conditions could help understand the rise of multidrug resistance in complex bacterial communities.

In summary, our results challenge the traditional cost/resistance trade off hypothesis, as functional ARCs can confer a growth benefit in the absence of selective pressure both *in vitro* and *in vivo*; and these effects are modulated by environmental conditions like the presence of oxygen. The joint structure of aerobic and anaerobic fitness, including how benefits and costs are distributed across environments, determines which ARCs can persist through unfavorable phases and expand when conditions improve. These findings indicate that ecological variability can sustain resistance cassettes that would otherwise be eliminated under constant conditions, and offer a broader mechanistic perspective on the stability of AMR determinants across heterogeneous environments.

## Methods

### Bacterial strains, plasmids, and culture conditions

*Escherichia coli* MG1655 was used as the host strain for all plasmid constructions in this study (Supplementary Table S7). Bacterial cultures were grown at 37°C in liquid Müller Hinton (MH; Oxoid, UK) or lysogeny broth (LB; Oxoid, UK) media, as well as on solid MH/LB agar (1.5%) plates (BD, France). To maintain the pMBA collection in *E. coli*, Zeocin (Invivogen, USA) was added at a concentration of 100 μg/mL. Liquid cultures were shaken at 200 rpm in an Infors Multitron shaker (Infors HT, Switzerland). The specific plasmids generated in this study, pMBA□ GFP KO and pMBA□ ΔGFP, were created by PCR amplification from corresponding pMBA derivatives, followed by Gibson Assembly, transformation, and Sanger sequencing. The oligonucleotides used for these procedures are listed in Supplementary Table S8.

### *In vitro* competition assays by flow cytometry

To evaluate the fitness cost of each ARC, we conducted *in vitro* competition assays following the protocol outlined in DeLaFuente et al., 2020^26^. Briefly, strains carrying different ARCs were competed against a strain containing the empty vector pMBA with a GFP knockout (GFP KO), allowing population differentiation via flow cytometry. By measuring the initial and final populations of each strain throughout the experiment, we determined the fitness cost associated with each ARC. Each competition was performed in six biological replicates per condition. Precultures were grown overnight in MH broth at 37°C with shaking, without selective pressure. A 1:1 mixture of the competing strains was prepared and diluted 1/400 in filtered saline solution to assess the initial population distribution. Measurements were taken using a Cytoflex-S flow cytometer (Beckman Coulter, US). The mixture was then diluted 1/400 in fresh MH broth and incubated at 37°C with shaking for 22 hours, after which the final proportions were quantified as before. The fitness of each pMBA derivative strain relative to the parental strain (pMBA KO) was calculated using the following formula:

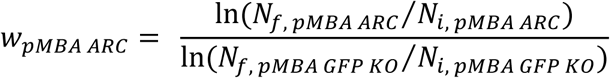

In the formula, w_pMBA ARC_ represents the relative fitness of each pMBA ARC in comparison with pMBA GFO KO. N_i_ and N_f_ corresponds to the number of cells at both the beginning and end of each competitor’s growth during the experiment. Data retrieved from this calculations was latter corrected using the formula achieved from the experimental regression line forced to y=1 in the intercept (y = 7.64*10^-7^x + 1), where x represents the cost and x the fluorescence intensity of the strain.

### OMM^12^ stocks preparation

The Oligo Mouse Microbiota (OMM^12^) is a defined bacterial community comprising twelve different bacterial species *(Enterococcus fecalis* KB1 (DSM32036), *Akkermansia muciniphila* YL44 (DSM26127), *Acutalibacter muris* KB18 (DSM26090), *Muribaculum intestinale* YL27 (DSM28989), *Flavonifractor plautii* YL31 (DSM26117), *Enterocloster clostridioformis* YL32 (DSM26114), *Bifidobacterium animalis* YL2 (DSM26074), *Turicimonas muris* YL45 (DSM26109), *Clostridium innocuum* I46 (DSM26113), *Bacteroides caecimuris* I48 (DSM26085), *Limosilactobacillus reuteri* I49 (DSM32035) and *Blautia coccoides* YL58 (DSM26115)). The OMM^12^ can colonize the murine intestinal tract and serve as a model for bacterial gut consortium in mice, as their members represent the five major eubacterial phyla found in the gut^28,49,50^.

OMM^12^ stocks for *in vitro* competition assays were prepared from individual monoculture inocula. These monocultures were grown from frozen glycerol stocks under strict anaerobic conditions at 37°C in 10 mL culture flasks (T25, Sarstedt, Germany) containing anaerobic medium (AMM) composed of 18 g/L glucose-free brain-heart infusion (Oxoid), 15 g/L glucose-free trypticase soy broth (USBiological), 5 g/L yeast extract, 2.5 g/L K₂HPO₄, 1 mg/L haemin, 0.5 mg/L menadione, 3% heat-inactivated fetal calf serum (FCS), and 0.25 g/L cysteine-HCl‧H₂O. The monocultures were then diluted 1:10 in fresh anaerobic medium and incubated for an additional 24 hours.

Before generating the OMM^12^ consortium, Gram staining was performed to check for contamination. Once confirmed to be contamination-free, monocultures were adjusted to an OD600 of 0.1 in fresh AMM medium and combined in equal proportions. The final OMM^12^ stock was aliquoted and stored at −80°C in anaerobic vials with glycerol at a final concentration of 10%. A 500 mL aliquot was collected to determine the relative abundance of each OMM^12^ member in the final mixture using qPCR.

### *In vivo* competition assays

*In vivo* competition assays were conducted using C57BL/6 mice stably colonized with the OMM^12^ defined bacterial consortium. The mice were housed under germ-free conditions in IsoCage P systems (Tecniplast, Italy) and provided with autoclaved ddH₂O and ad libitum access to mouse-breeding complete food (Ssniff, Germany). All mice used were males aged 10–15 weeks, randomly assigned to experimental groups, and housed in groups of 2–4 per cage. Animal health was monitored twice daily throughout the experiment.

Prior to bacterial inoculation, strains were cultured in MH broth at 37°C for 12 hours on a wheel rotor and mixed at a 1:1 ratio (pMBA_ARC_ ΔGFP vs. pMBA□ ΔGFP). Mice were inoculated via gavage with the bacterial mixture (100 µL applied to the fur and 50 µL administered orally). Fecal samples were collected every 24 hours for the first four days and on day 7, the final day of the experiment, for plating and quantitative PCR (qPCR) analysis. All mice were euthanized by cervical dislocation on day 7, and their cecal contents were collected.

*E. coli* loads in feces and cecal contents were quantitated by plating on MacConkey agar (Oxoid) supplemented with vancomycin (7.5 mg/mL). 100 colonies were restreaked on MacConkey agar with both vancomycin and zeocin to determine *E. coli* pMBA loads. The proportion of resistant colonies was inferred by replica plating 50 colonies per sample onto MacConkey agar plates with selective conditions specific to the ARC encoded in the vector: *ereA2* (erythromycin 250 μg/mL), *aacA7* (kanamycin 25 μg/mL), and *bla*_OXA-10_ (carbenicillin 100 μg/mL).

### *In vitro* competition assays by phenotypic selection

Bacterial fitness in the presence of each ARC was further evaluated by phenotypic selection across various environments and conditions. In these experiments, pMBA-derived strains without GFP (pMBA_ARC_ ΔGFP) were competed against a strain containing the empty vector without GFP (pMBA□ ΔGFP). At least three independent biological replicates were conducted for each competition and condition. We carried out three separate competition experiments, selecting by phenotype in the following environments: with oxygen, without oxygen, and in the presence of OMM^12^ bacterial consortium under anaerobic conditions.

Aerobic competitions were carried out at 37°C with shaking. Precultures of both pMBA_ARC_ ΔGFP and pMBA□ ΔGFP were grown overnight in MH broth. Then, 10⁶ CFUs from each culture were mixed in 5 mL of MH broth without selective pressure and incubated overnight. Every 24 hours for 5 days, 2 × 10^6^ CFUs were transferred into 5 mL of fresh MH broth. Samples were taken daily, diluted 100,000 times, and plated onto MH agar plates. The proportion of resistant colonies was determined by replica plating 50 colonies per sample onto MH agar plates with selective pressure specific to the ARC encoded in the vector: *ereA2* (erythromycin 250 μg/mL), *dfrA21* and *dfrA31* (trimethoprim 50 250 μg/mL), *aacA7* (kanamycin 25 μg/mL), and *bla*_OXA-10_ (carbenicillin 100 μg/mL).

Anaerobic competitions were conducted in a controlled gas atmosphere (7% H₂, 10% CO₂, 83% N₂). Precultures were grown overnight in 24-well plates (Nunc) and mixed by adding 0.1 mL of each culture (OD600 = 0.1) to a final volume of 1 mL of fresh MH broth. After 24 hours of incubation, the mixtures were diluted 1:100 in fresh MH every 24 hours for 5 days. Plating was done as described in the previous section.

Anaerobic competitions with OMM^12^ consortium followed the same protocol, except the initial proportions in the mixture were 10:1. A total of 100 µL of OMM12 stock and 10 µL of each competitor were added to 900 µL of fresh M10 broth (18 g/L glucose-free brain-heart infusion (BHI; Oxoid), 15 g/L glucose-free trypticase soy broth (TSB; USBiological, USA), 5 g/L yeast extract, 2.5 g/L K₂HPO₄, 1 mg/L haemin, 0.5 mg/L menadione, 3% heat-inactivated fetal calf serum, 0.25 g/L cysteine-HCl·H₂O, 0.025% mucin, 0.5 g/L of fucose, xylose, arabinose, rhamnose, lyxose, and 2 g/L of xylan from beechwood (Roth, Germany)). Additionally, 500 mL of the mixed cultures on day 5 were collected to quantify the relative abundance of each OMM^12^ member.

### gDNA extraction from fecal pellets, cecal samples and *in vitro* competitions

We performed genomic DNA (gDNA) extraction from competitions were OMM^12^ community was present in order to quantitate the relative abundance of each member. The protocol used was stablished based on *Turnbaugh et at., 2009* and *Ubeda et al., 2012* as references^51,52^.

Fecal, cecal or *in vitro* competition samples were thawed and resuspended in 500 mL of extraction buffer (20 mM EDTA, 200 mM NaCl, 200 mM Tris (pH 8.0)), 210 mL of 20% SDS, 500 ml phenol:chloroform:isoamylalcohol (25:24:1, pH 7.9, Roth) and 500 mL of 0.1 mm-diameter zirconia/silica beads (BioSpec Products, Bartlesville, OK, Roth). Bacteria were lysed using a bead beater (TissueLyser LT, Qiagen) for 4 minutes at max speed (50 s^-1^), followed by centrifugation at full speed (14,000 x g) for 5 min. Supernatants were collected avoiding the phenol from the lower phase and mixed with 500 mL of phenol:chloroform:isoamylalcohol. Samples were centrifuged again, collecting the upper phase of the supernatant. To precipitate gDNA, this phase was then mixed by inversion with 1 mL of 96 % ethanol supplemented with 50 ml Sodium Acetate 3M. After 30 min centrifugation (14,000 x g), at 4 °C; pellets were washed with 500 ml 70% ice-cold ethanol and centrifuged again 15 min (14,000 x g), at 4 °C. The gDNA pellets were air-dried and then resuspended in 100 µL of TE buffer. Additionally, the gDNA samples were purified using the NucleoSpin gDNA clean-up kit (Macherey-Nagel, Germany), according to the manufacturer’s guidelines.

### Quantification of OMM^12^ microbiota members and *E. coli* abundance by qPCR

To assess the relative abundance of OMM^12^ members and *E. coli* during in competition assays, we performed qPCR of bacterial 16S rRNA genes according to *Brugiroux et al, 2016* protocol^49^. Prior to the experiments, qPCR reactions were optimized to achieve efficiencies between 90-110%. qPCR assays using gDNA extracted were performed in duplicates using a LightCycler96 Thermocycler (Roche, Switzerland). Each reaction contained 30 μM of each primer, 25 μM of each probe, FastStart Essential DNA Probes MasterMix (Roche), and 5 ng of template gDNA. The thermocycling conditions included an initial 10-minute denaturation at 95°C, followed by 45 cycles of amplification of 15 seconds at 95°C and 1 minute at 60°C. Fluorescence was recorded after the 60°C step for each cycle. The quantification cycle (Cq) and baseline were automatically determined using the LightCycler96 software version 1.1 (Roche). The primers and probes used for are listed in Supplementary Table S8.

### Growth curves

Growth curves shown in Supplementary Figure S1 were performed as follows: Three independent bacterial colonies were incubated overnight at 37°C with shaking in MH medium containing zeocin. The cultures were then diluted 1:200 in fresh MH zeocin medium, and 200 µL of the dilution were inoculated into 96-well plates. The plates were incubated for 22 hours, measuring OD600 of each well every 10 minutes using an incubated Synergy HTX plate reader. Before each measurement, the plates were shaken at 280 cycles per minute.

### Statistical analysis

Statistical analyses were performed using GraphPad Prism version 10 with the exception of estimations retrieved from a linear regression model. Specific statistical tests used for each dataset are described in figure and table legends. To assess if the insertion of the gene had an impact on bacterial fitness, measures of fitness effect with and without each gene were analyzed using linear regression models separately for aerobic and anaerobic conditions considering first the effect of the antimicrobial family and then the individual gene. Briefly, the fitness measure was used as a response variable for lineal models including aerobic/anaerobic conditions (as a dichotomous variable) and the gene family or the specific gene (alternatively) as covariates, so that the coefficients associated with each covariate measured the variation above/below the baseline condition.

### Stochastic population dynamics model

We used a stochastic, resource-explicit competition model to simulate the dynamics of ARC-bearing populations. Each simulation represents a pairwise competition between one ARC-bearing strain and the isogenic reference strain carrying pMBA∅. Population growth follows Monod kinetics, where the instantaneous division rate of strain *i* is

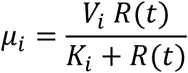

with *V_i_* the maximum growth rate, *K_i_* the half-saturation constant, and *R(t)* the concentration of a single limiting resource. Resource uptake is proportional to biomass production, and resource depletion is tracked continuously. Populations are propagated through daily growth-dilution cycles, similarly to the *in vitro* protocol, with a 1:100 transfer between cycles. An ARC is considered extinct when its population falls below one cell at the end of a cycle.

Simulations were performed under three environmental regimes: fully aerobic, fully anaerobic, or a single-switch regime where cultures grow anaerobically for *K* days before switching permanently to aerobiosis. Simulations were initialized with equal densities of the ARC and control strain (1:1 ratio), and a fixed initial resource concentration. Each strain has its own set of growth parameters *(Vi, Ki)*, estimated by mapping experimentally measured relative fitness to condition-specific growth parameters using growth-rate measurements and short-term competition assays, following the calibration procedure described in the Supplementary Information. To represent the empirical variation within each ARC family, we fitted a bivariate aerobic–anaerobic fitness distribution (family-specific means, variances, and covariance) and used this joint distribution to generate synthetic competitors. For each family, the resulting library included all experimentally measured cassettes and simulated genotypes drawn from the same distribution. Synthetic genotypes therefore preserve not only the marginal fitness variation but also the covariance structure observed across environments.

For every environmental regime and genotype, we performed multiple stochastic replicates (1,000 per family). At each transfer we recorded population sizes, frequencies, and extinction events. Persistence probability was defined as the fraction of replicates in which the ARC population remained above the extinction threshold after 100 cycles. Rescue was defined as persistence under a switching regime for genotypes that went extinct under continuous anaerobiosis. All simulations were implemented in Python using a fixed time-step tau-leaping scheme, where birth events were treated as Poisson processes with propensities defined by the Monod growth terms. The full implementation of the model, together with the data and scripts used to generate all simulations, is available at: https://github.com/ccg-esb-lab/ARCfitness

**Figures:** Most figures were created with BioRender.com.

## Supporting information

Mathematical model

## Data availability

All data reported in the manuscript are either represented in the figures or in Supplementary Material. Source data are provided with this paper.

## Acknowledgements

We would like to thank all the members of MBA lab and Stecheŕs lab for discussions, especially Laura Ortiz and Amalia Prieto. The work in the MBA laboratory is supported by the European Research Council (ERC) through a Starting Grant [ERC StG grant no. 803375-KRYPTONINT] and the Ministerio de Ciencia e Innovación [PID2020-117499RB-I00 and CNS2022-135857]; and the EU HARISSA JPI-AMR program [PCI2021-122024-2A]. A.H. has also been supported by grants from EMBO scientific exchange grant number 9504 and FEMS research and training grant ID FEMS-GO-2021-084. R.P.-M. and A.F.-H. were supported by a DGAPA-PASPA 2024 sabbatical grant from UNAM. B.S. received funding by the European Research Council (ERC) under the European Union’s Horizon 2020 research and innovation program (grant agreement 865615), the German Research Foundation (DFG, German Research Foundation, SFB 1371, project number 395357507), and the German Center for Infection Research (DZIF). Work in the A.S.M. lab was supported by the European Research Council (ERC) under the European Union’s Horizon 2020 research and innovation programme (ERC StG grant no. 757440-PLASREVOLUTION)

## Author contributions

Conceptualization: JAE, AH.

Investigation: AH, LGP, AS, AW, AF-H, RP-M, JSD, EV, JA.

Funding acquisition: JAE, BS, ASM.

Project administration: JAE.

Supervision: JAE, BS, ASM.

Writing Original Draft: AH, JAE.

Writing review and editing: AH, JAE, BS, ASM.

## Ethical

Germ-free and OMM^12^- colonized C57BL/6J mice were bred and maintained in flexible film isolators (North Kent Plastic Limited, U.K.) at germ-free mouse facilities at the Max-von-Pettenkofer Institute (Facility MUC; Munich, Germany). All animal experiments were reviewed and approved by the local authorities (Regierung von Oberbayern, under the ethical approval number ROB-55.2-2532.Vet_02-20-118).

