## Supplementary Material for "Fitness effects of antimicrobial resistance genes in changing environments"

### pMBA$_{ARC}$ GROWTH CURVES

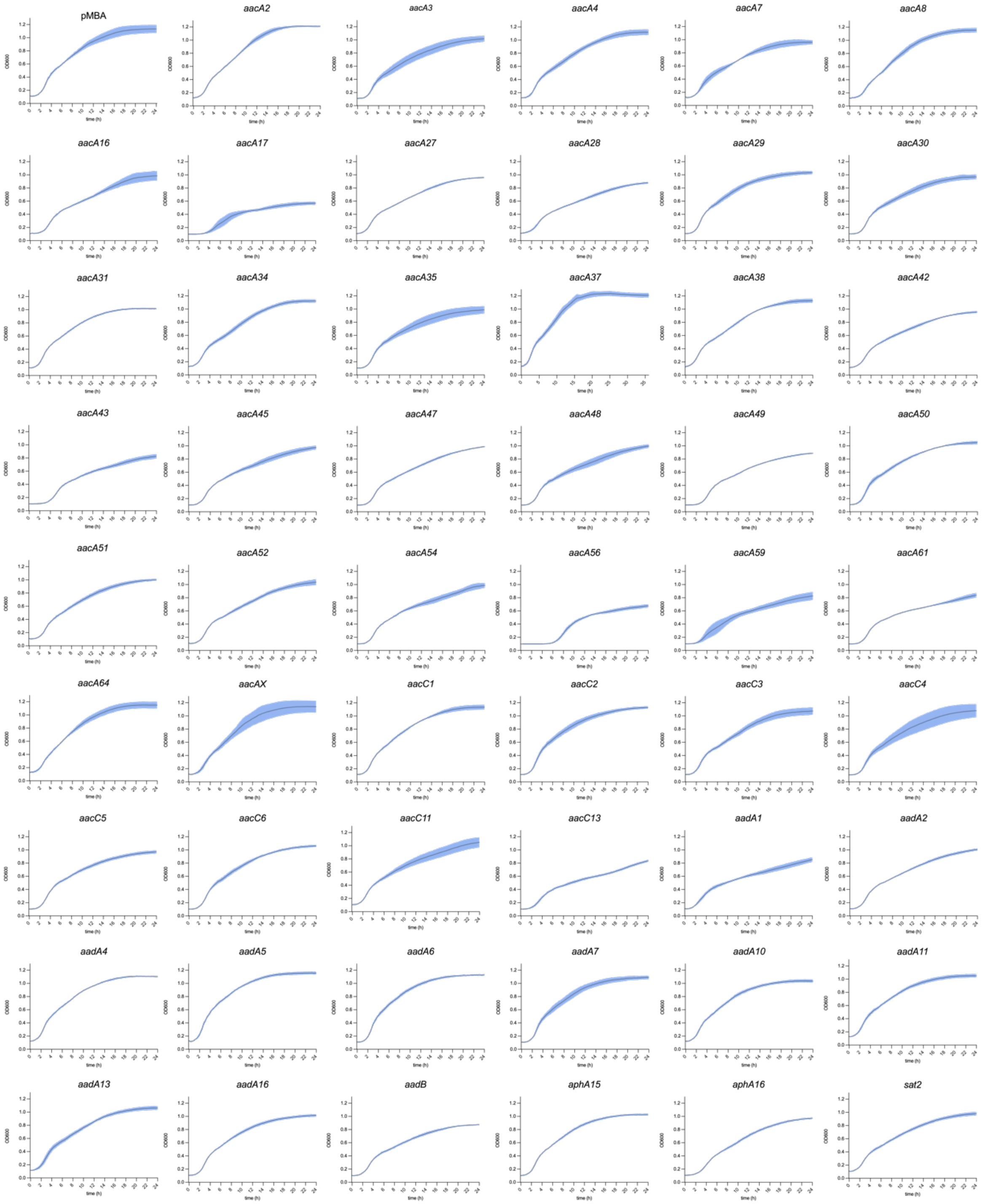

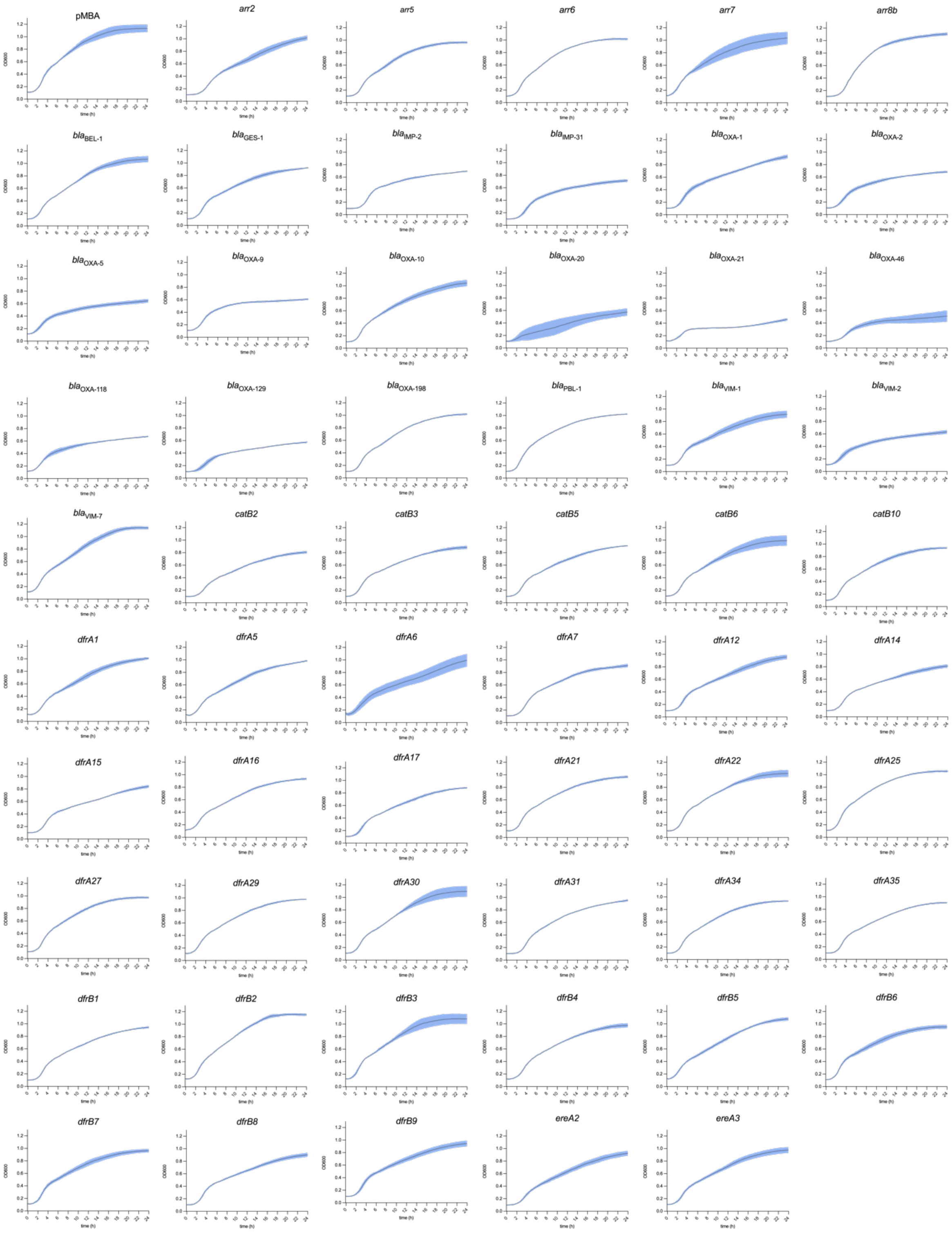
pMBA
arr2
arr5
arr6
arr7
arr8b
$bla_{BEL-1}$
$bla_{GES-1}$
$bla_{IMP-2}$
$bla_{IMP-31}$
$bla_{OXA-1}$
$bla_{OXA-2}$
$bla_{OXA-5}$
$bla_{OXA-9}$
$bla_{OXA-10}$
$bla_{OXA-20}$
$bla_{OXA-21}$
$bla_{OXA-46}$
$bla_{OXA-118}$
$bla_{OXA-129}$
$bla_{OXA-198}$
$bla_{PBL-1}$
$bla_{VIM-1}$
$bla_{VIM-2}$
$bla_{VIM-7}$
catB2
catB3
catB5
catB6
catB10
dfrA1
dfrA5
dfrA6
dfrA7
dfrA12
dfrA14
dfrA15
dfrA16
dfrA17
dfrA21
dfrA22
dfrA25
dfrA27
dfrA29
dfrA30
dfrA31
dfrA34
dfrA35
dfrB1
dfrB2
dfrB3
dfrB4
dfrB5
dfrB6
dfrB7
dfrB8
dfrB9
ereA2
ereA3
OD600
time (h)

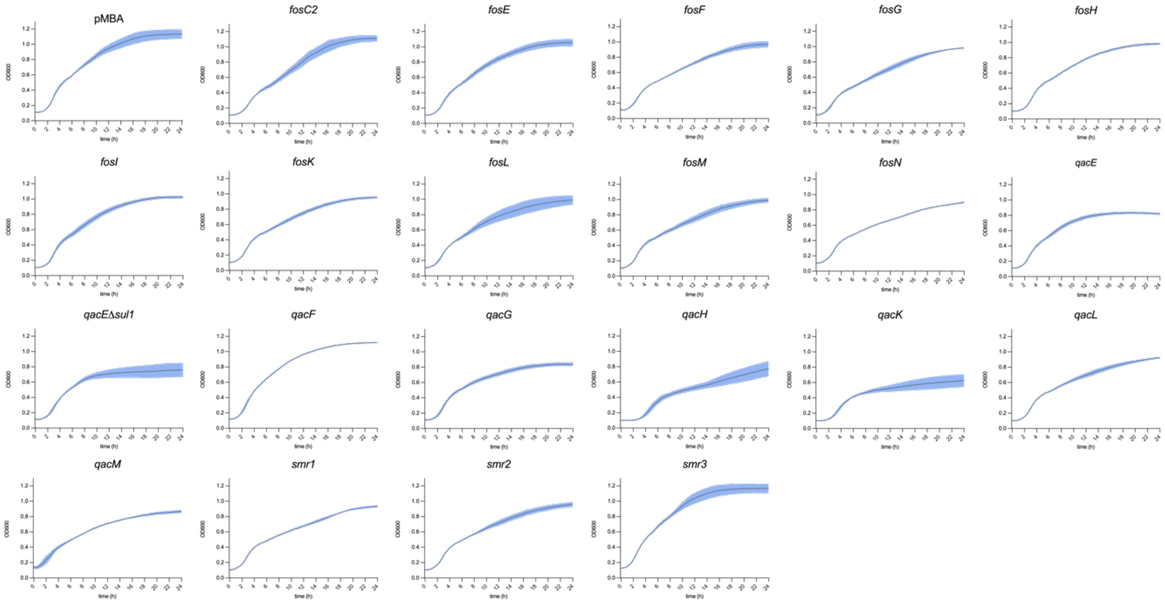


**Supplementary Figure S1. Growth curves of all pMBA$_{ARCs}$.** Growth of each pMBA derivate was assessed along 24h as described in materials and methods. Graphs represent the mean of three independent replicates being the standard error of the mean (SEM) represented as a shadow in lighter colour.

A

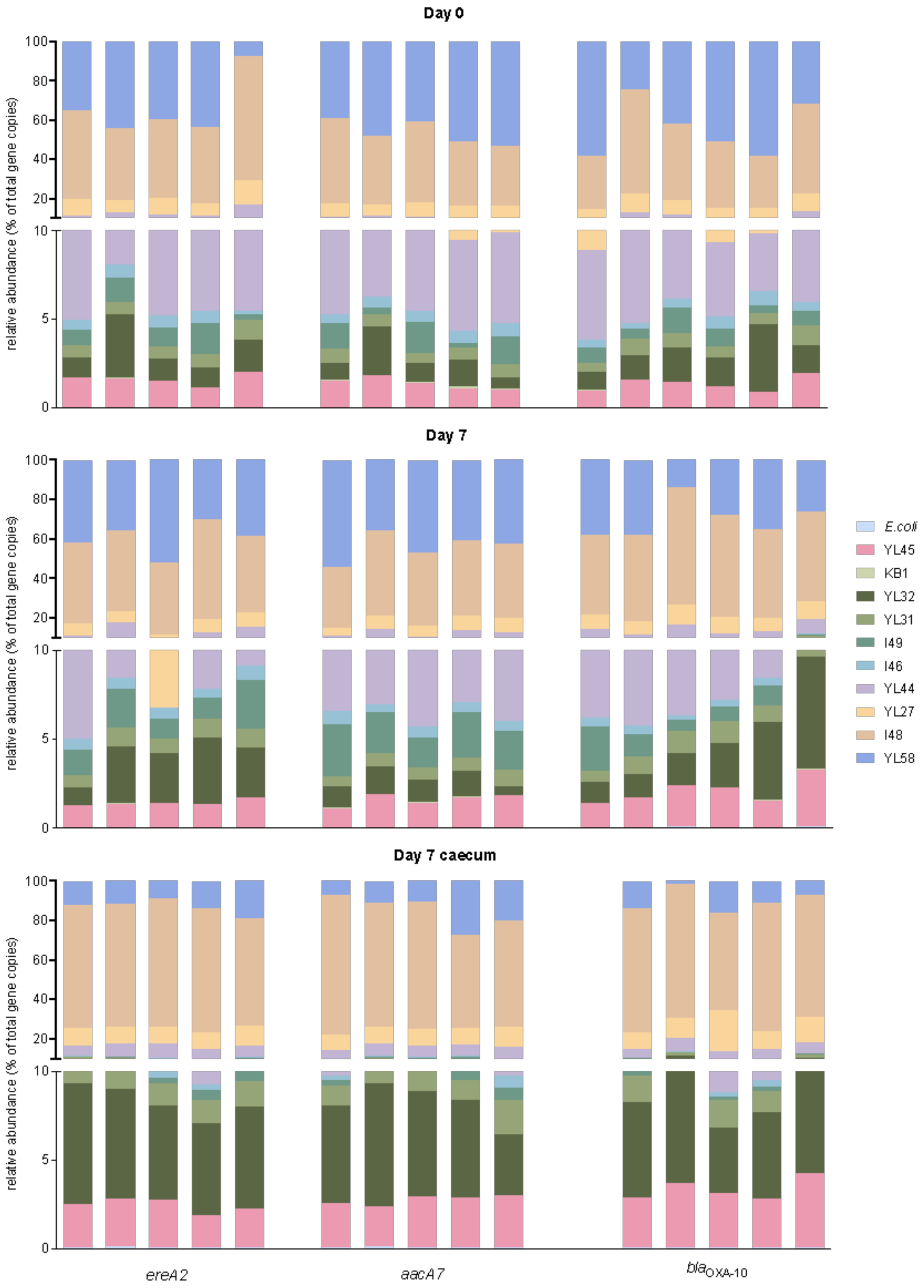
Day 0
Day 7
Day 7 caecum
relative abundance (% of total gene copies)
100
80
60
40
20
10
5
0
ereA2
aacA7
blaOXA-10
E.coli
YL45
KB1
YL32
YL31
I49
I46
YL44
YL27
I48
YL58

B

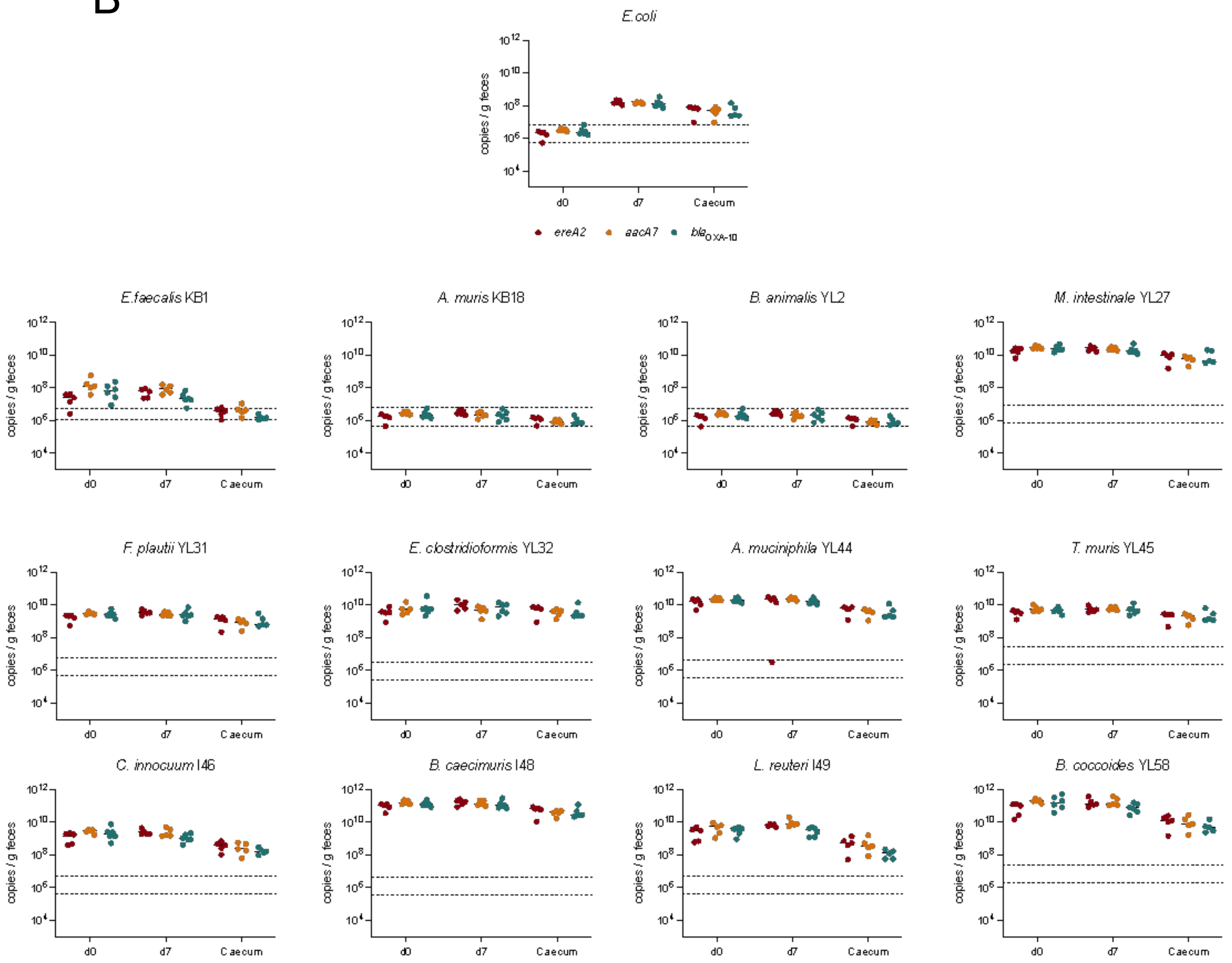


**Supplementary Figure S2. Quantitation of OMM$^{12}$ members and *E. coli* abundance during *in vivo* competitions. A)** Relative abundance in mice before (day 0) and after (day 7 and caecum) *in vivo* competitions. YL2 and KB18 are not present in the graph as their levels are under the limit of quantification**.** *E. coli* and OMM$^{12}$ members are represented in different colors following the figure legend pattern. **B)** Absolute quantification of bacterial species measured by qPCR (copies/ g feces). Dotted lines represent de limit of quantification and the limit of detection of the technique. Each competition is represented with a different color: red (*ereA2*), yellow (*aacA7*) and blue (*bla*$_{OXA-10}$). Data is represented as the median of at least 5 individuals.

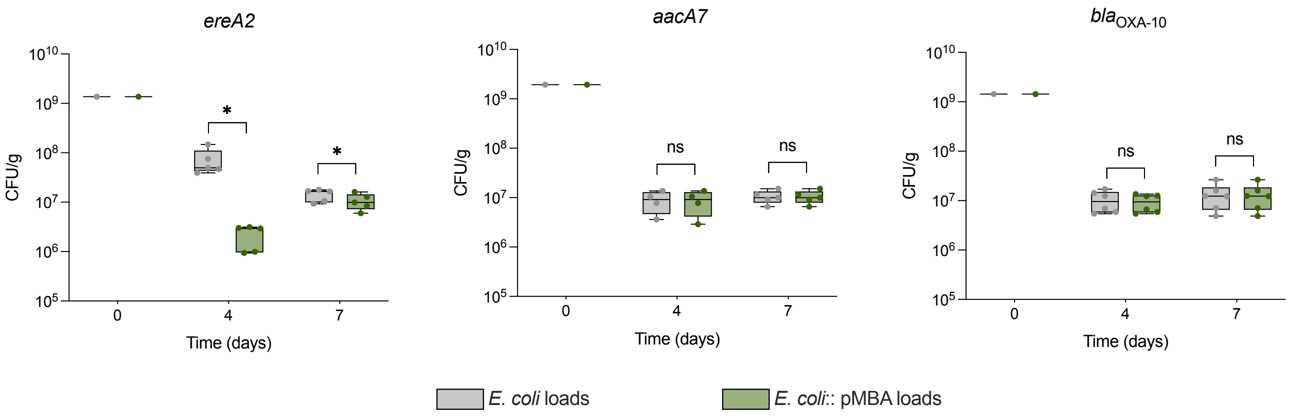


**Supplementary Figure S3. Quantification of plasmid loss during *in vivo* experiments.** Quantitation of total *E. coli* loads (grey boxes) and pMBA-containing *E. coli* (green boxes) at days 0, 4 and 7 of *in vivo* competitions pMBAØ vs pMBA$_{ereA2}$, pMBA$_{aacA7}$ or pMBA$_{bla\text{OXA-10}}$ (from left to right). Data is represented using a box-and-whisker plot, where each experimental group consists of at least five individuals. Statistical differences were assessed using a two-sided pair t-test. * P < 0.05. ns = non significative.

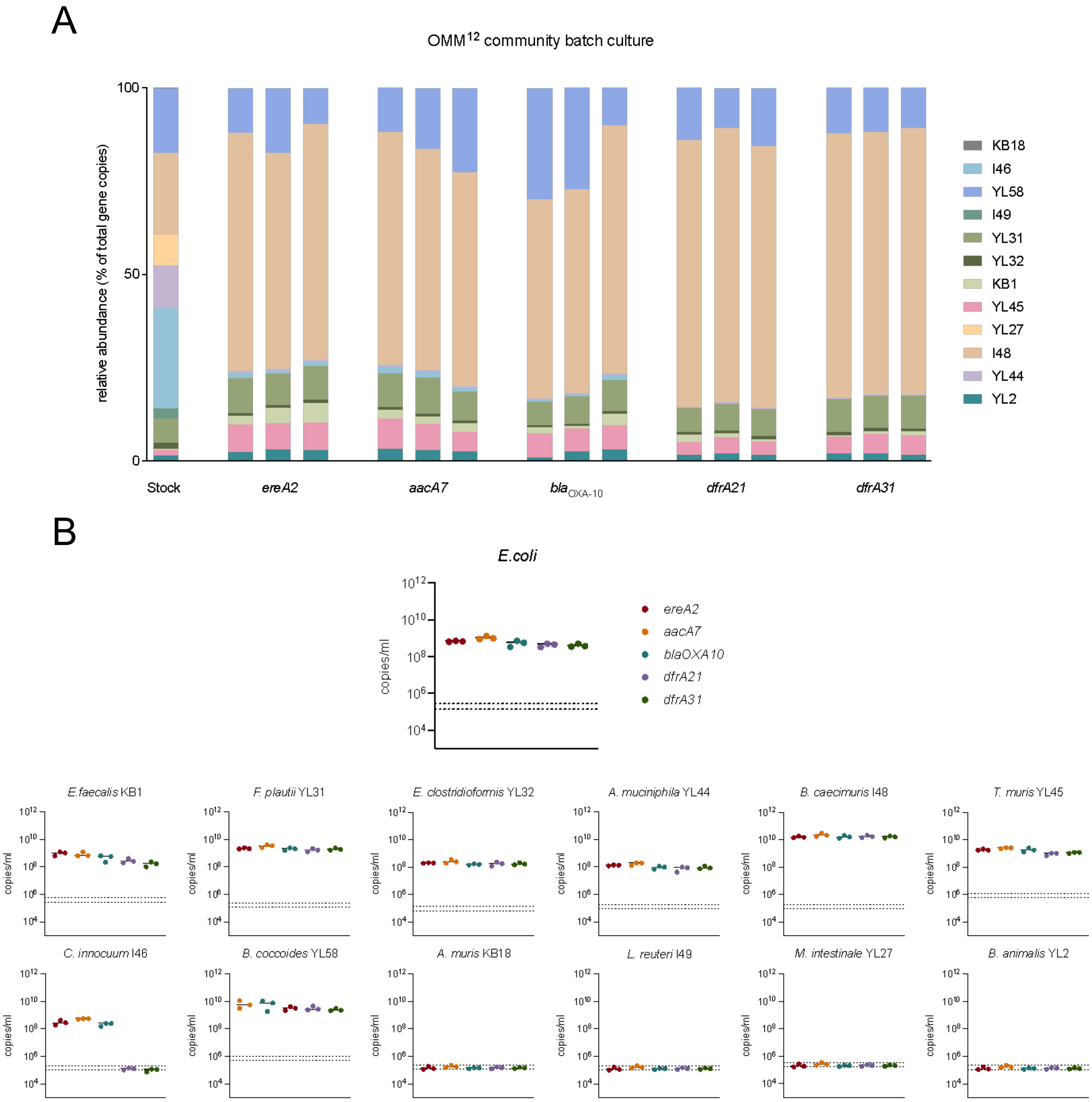


**Supplementary Figure S4. Quantitation of OMM$^{12}$ members and *E. coli* abundance during *in vitro* anaerobic long -term competitions. A)** Relative abundance of OMM$^{12}$ community stock (day 0) and after *in vitro* competitions (day 4). OMM$^{12}$ members are represented in different colors following the figure legend pattern. **B)** Absolute quantification of bacterial species measured by qPCR (copies/ mL). Dotted lines represent de limit of quantification and the limit of detection of the technique. Each competition is represented with a different color: red (*ereA2*), yellow (*aacA7*) and blue (*bla*$_{OXA-10}$), purple (*dfrA21*) and green (*drfA31*). Data is represented as the median of 3 biological replicates.

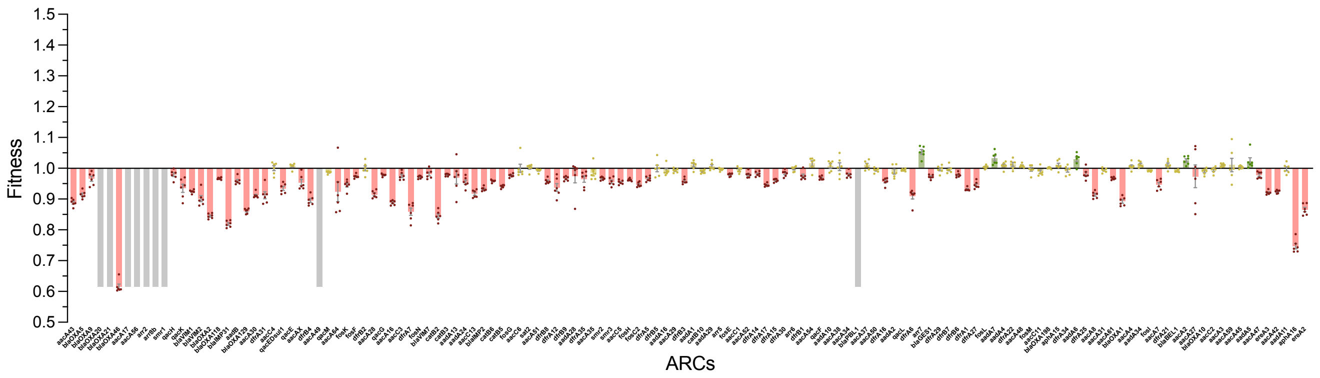


**Supplementary Figure S5. ARC fitness effect characterization in the absence of oxygen.** Graph showing the fitness effect of each ARC on bacterial growth in anaerobic conditions. ARCs are ordered by their fitness effect in aerobic conditions. Bars represent the mean of 6 independent replicates, error bars represent the standard error of the mean (SEM). Grey bars represent ARCs unable to be measured, red bars costly ARCs, yellow bars represent ARCs with no significant cost (according to a linear regression model), and green bars state for beneficial ARCs under no selective pressure.

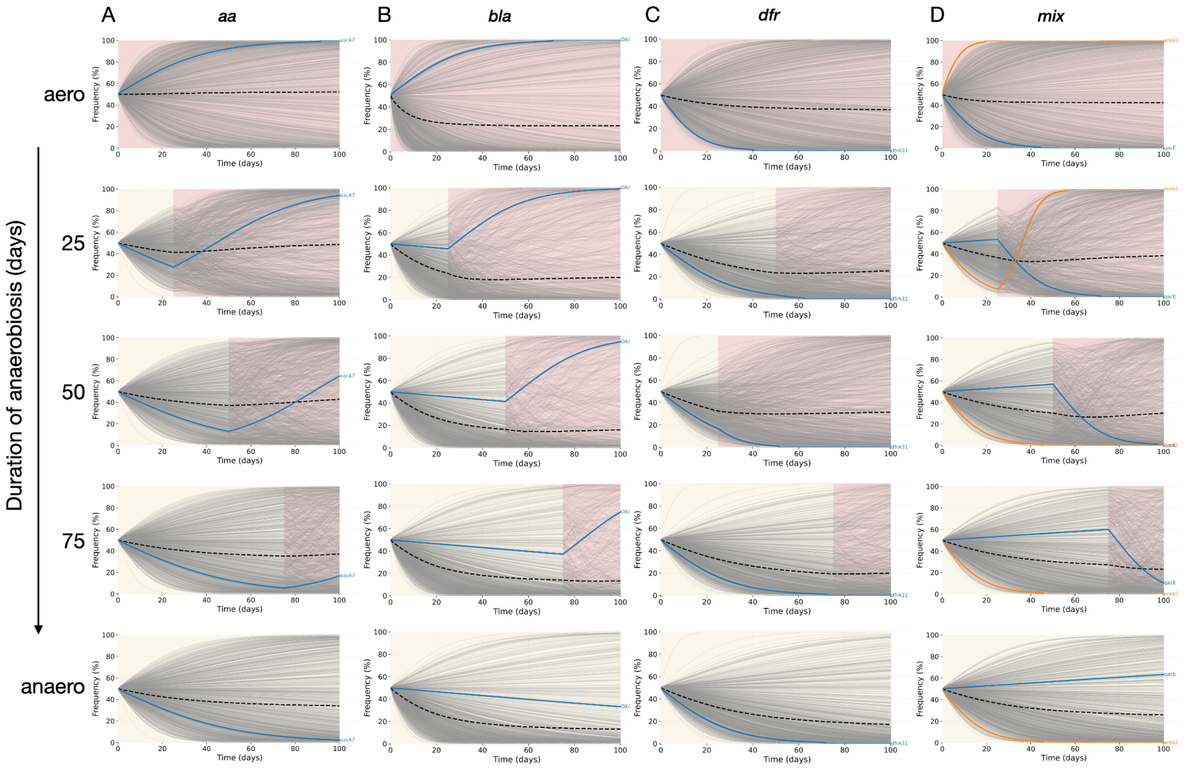


**Supplementary Figure S6.** Long-term competition dynamics across families under increasing durations of anaerobic exposure. Each column corresponds to an ARC family (*aa*, *bla*, *dfr*, and *mix*), and each row shows simulations under a different duration of anaerobiosis (0, 25, 50, 75, and 100 days). Panels display the time-resolved frequency of the ARC-bearing strain during daily transfers. Light grey lines show trajectories from the synthetic fitness library, the dotted black line marks the family-level mean trajectory, and colored lines highlight experimentally measured ARCs included in the simulations. Background shading indicates the environmental state: yellow for anaerobic conditions and red for aerobic conditions.

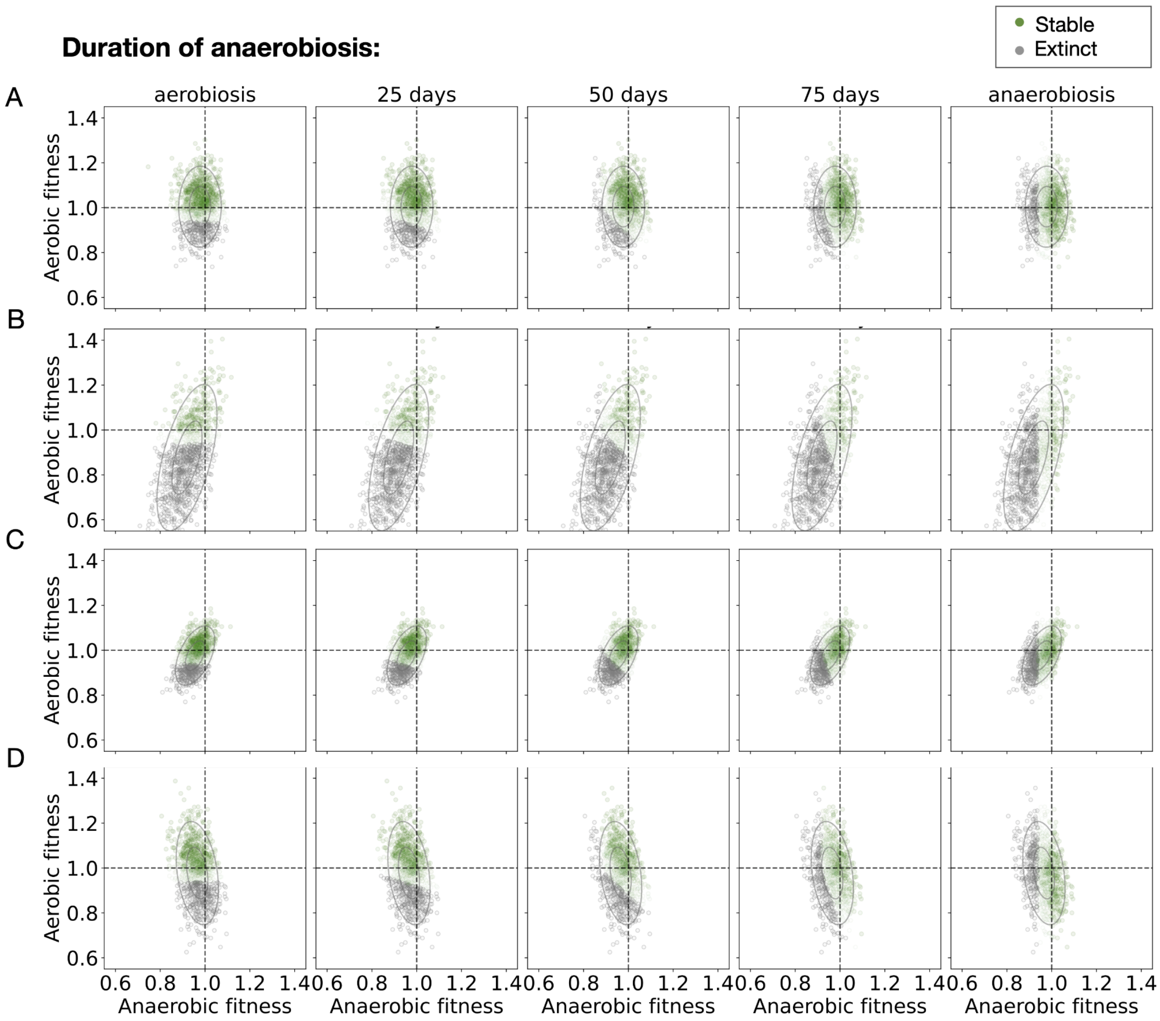


**Supplementary Figure S7. Joint fitness landscapes under increasing durations of anaerobiosis.** Matrix of joint fitness landscapes showing aerobic and anaerobic fitness for all simulated ARC-bearing strains. Rows correspond to ARC families (top to bottom: *dfr, bla, aa, mix*), and columns reflect increasing durations of the initial anaerobic phase before switching to aerobiosis. Each point represents an individual ARC, with green indicating strains that persisted after 100 simulated transfers and gray those that went extinct. Under constant aerobiosis (leftmost column), survival is primarily linked to high aerobic fitness. Under constant anaerobiosis (rightmost column), survival depends on anaerobic fitness. In intermediate conditions, the survival boundary gradually rotates from vertical to horizontal, forming a rescue zone in the upper-right quadrant of the landscape. This pattern shows that the anaerobic-to-aerobic transition selects for asymmetric fitness profiles across environments. Strains with high fitness in one condition and lower fitness in the other fall within the rotated survival region and can be rescued by the switch, whereas those with low fitness in both environments remain outside this region and go extinct.

**Supplementary Table 1. Fitness of ARCs.** Using a linear regression model we estimate the coefficient of variation in the fitness of the host due to presence of the ARC and its significance (p-value). *qacEDsul1* was excluded of the analysis.

| ARC | Mean Fitness | Estimate Coefficient | p.value | Signif. |
|---|---|---|---|---|
| *aacA16* | 0,89103033 | -0.108969665915153 | 2.54383450158282e-09 | Yes |
| *aacA2* | 1,09047811 | 0.0904781099114083 | 6.70805464470306e-07 | Yes |
| *aacA27* | 1,0940067 | 0.0940066961825542 | 2.4781288515582e-07 | Yes |
| *aacA28* | 0,88969898 | -0.110301020494876 | 1.64671472892138e-09 | Yes |
| *aacA29* | 0,97797824 | -0.0220217583877921 | 0.222016882548585 | No |
| *aacA3* | 1,09970605 | 0.0997060486094533 | 4.63406467646282e-08 | Yes |
| *aacA30* | 0,84224679 | -0.157753208072881 | 1.97023449562901e-17 | Yes |
| *aacA31* | 1,06641694 | 0.0664169410715887 | 0.000247319531029451 | Yes |
| *aacA34* | 1,07292802 | 0.0729280246942136 | 5.82478373829894e-05 | Yes |
| *aacA35* | 0,96626816 | -0.0337318372379544 | 0.061621311235983 | No |
| *aacA37* | 1,01302322 | 0.013023224071586 | 0.470005551463561 | No |
| *aacA38* | 1,00381505 | 0.00381504506736715 | 0.832354351366391 | No |
| *aacA4* | 1,07173937 | 0.0717393734847932 | 7.64960652396299e-05 | Yes |
| *aacA42* | 1,15057275 | 0.150572748262241 | 4.31102114466878e-16 | Yes |
| *aacA45* | 1,10215454 | 0.102154536493298 | 2.19786256289945e-08 | Yes |
| *aacA47* | 1,1141063 | 0.114106296802207 | 4.63704085310669e-10 | Yes |
| *aacA48* | 1,03298359 | 0.0329835862156377 | 0.0675996281489038 | No |
| *aacA49* | 0,87276956 | -0.127230439681827 | 4.47057965122615e-12 | Yes |
| *aacA50* | 1,01374367 | 0.0137436718201931 | 0.445810163303787 | No |
| *aacA51* | 0,94818472 | -0.051815284284993 | 0.00416457840393647 | Yes |
| *aacA52* | 0,99213752 | -0.00786248392237139 | 0.662666485138717 | No |
| *aacA54* | 0,99780438 | -0.00219562193379001 | 0.903034773263333 | No |
| *aacA59* | 1,1017635 | 0.101763503453451 | 2.47847590695871e-08 | Yes |
| *aacA61* | 1,06718645 | 0.0671864498943634 | 0.000209720371773661 | Yes |
| *aacA64* | 0,88137021 | -0.118629790735572 | 9.81408382895727e-11 | Yes |
| *aacA7* | 1,07385231 | 0.0738523114783488 | 4.70002592552299e-05 | Yes |
| *aacA8* | 1,06611206 | 0.0661120565962777 | 0.00026390081683487 | Yes |
| *aacAX* | 0,86763853 | -0.13236147387599 | 6.51363448178165e-13 | Yes |
| *aacC1* | 0,98673099 | -0.0132690053788976 | 0.46167053579449 | No |
| *aacC11* | 1,03721139 | 0.0372113875572908 | 0.0392871798852348 | Yes |
| *aacC13* | 0,9224326 | -0.0775673976664901 | 1.93865409385325e-05 | Yes |
| *aacC2* | 1,09839842 | 0.0983984184578898 | 6.85872941971349e-08 | Yes |
| *aacC3* | 0,8974388 | -0.102561199696073 | 1.93888028898786e-08 | Yes |
| *aacC4* | 0,85980647 | -0.140193526871609 | 3.06236013986103e-14 | Yes |
| *aacC5* | 0,9668523 | -0.0331476979978856 | 0.0662490191092551 | No |
| *aacC6* | 0,94539829 | -0.0546017141133318 | 0.00254118855297353 | Yes |
| *aadA1* | 0,97981395 | -0.0201860525206634 | 0.26292872403659 | No |
| *aadA10* | 1,00251344 | 0.00251344082630935 | 0.889084167908812 | No |
| *aadA11* | 1,15357713 | 0.153577128769275 | 1.20132745249857e-16 | Yes |
| *aadA13* | 0,91682104 | -0.0831789649444479 | 4.74384379238295e-06 | Yes |
| *aadA16* | 0,97381803 | -0.0261819690827634 | 0.146636788185361 | No |
| *aadA2* | 1,01942629 | 0.0194262856024843 | 0.281301264950391 | No |
| *aadA24* | 0,92078123 | -0.0792187679628855 | 1.29234879739739e-05 | Yes |
| *aadA28* | 0,96074365 | -0.0392563455538057 | 0.0297042840177138 | Yes |
| *aadA29* | 0,98291783 | -0.0170821660885645 | 0.343386174432968 | No |
| *aadA34* | 1,00590273 | 0.00590273269723183 | 0.743280049408797 | No |
| *aadA4* | 1,03122263 | 0.0312226319287902 | 0.0835703586054944 | No |
| *aadA5* | 1,10684998 | 0.106849982721838 | 5.03754361958561e-09 | Yes |
| *aadA6* | 1,05102907 | 0.0510290659660828 | 0.00476916233393706 | Yes |
| *aadA7* | 1,03019667 | 0.0301966749927436 | 0.0942072126083767 | No |
| *aadB* | 0,78456162 | -0.215438384319576 | 9.26387344934761e-30 | Yes |
| *aphA15* | 1,04319276 | 0.043192760093347 | 0.0168008886711324 | Yes |
| *aphA16* | 1,18290779 | 0.18290778919624 | 1.74529994296036e-22 | Yes |
| *arr5* | 0,98354495 | -0.0164550456700854 | 0.361381622324869 | No |
| *arr6* | 0,99709859 | -0.00290141418452826 | 0.872101129570853 | No |
| *arr7* | 1,0216258 | 0.0216257979307883 | 0.230430220436606 | No |
| *bla*$_{BEL-1}$ | 1,07946355 | 0.0794635459171045 | 1.21619761774725e-05 | Yes |
| *bla*$_{GES-1}$ | 1,02206358 | 0.0220635792242336 | 0.221141323490614 | No |
| *bla*$_{IMP-2}$ | 0,92804086 | -0.0719591416900312 | 7.27579816907112e-05 | Yes |
| *bla*$_{IMP-31}$ | 0,73311741 | -0.266882586783711 | 1.22025404989602e-42 | Yes |
| *bla*$_{OXA-1}$ | 1,06758593 | 0.067585931805878 | 0.000192391475894046 | Yes |

| ARC | Mean Fitness | Estimate Coefficient | p.value | Signif. |
|---|---|---|---|---|
| *bla*$_{OXA-10}$ | 1,0972842 | 0.0972842045464649 | 9.54621415224918e-08 | Yes |
| *bla*$_{OXA-118}$ | 0,73300187 | -0.266998127680932 | 1.13753277229161e-42 | Yes |
| *bla*$_{OXA-129}$ | 0,78992238 | -0.210077623967472 | 1.65110220565007e-28 | Yes |
| *bla*$_{OXA-198}$ | 1,03982063 | 0.0398206347073672 | 0.027444401263399 | Yes |
| *bla*$_{OXA-2}$ | 0,73145378 | -0.268546218524624 | 4.43509830701568e-43 | Yes |
| *bla*$_{PBL-1}$ | 1,01032073 | 0.0103207302038903 | 0.566920047775218 | No |
| *bla*$_{VIM-1}$ | 0,69170197 | -0.308298028993115 | 6.4700031664572e-54 | Yes |
| *bla*$_{VIM-2}$ | 0,71342666 | -0.28657333780122 | 6.4205512614864e-48 | Yes |
| *bla*$_{VIM-7}$ | 0,91032628 | -0.0896737159302188 | 8.37927641485098e-07 | Yes |
| *catB10* | 0,98075113 | -0.0192488732934684 | 0.285714280945808 | No |
| *catB2* | 0,91085542 | -0.089144577286437 | 9.69070897663582e-07 | Yes |
| *catB3* | 0,91378134 | -0.0862186561217764 | 2.13682229039416e-06 | Yes |
| *catB5* | 0,94040188 | -0.0595981223793114 | 0.000993874323923482 | Yes |
| *catB6* | 0,94039647 | -0.0596035282306071 | 0.000992828759119365 | Yes |
| *dfrA1* | 1,02593279 | 0.0259327926350017 | 0.15051258043648 | No |
| *dfrA12* | 0,95459133 | -0.0454086737443989 | 0.0119680335475136 | Yes |
| *dfrA14* | 0,99228655 | -0.00771345432055794 | 0.668674932847416 | No |
| *dfrA15* | 0,99533522 | -0.004664777838243 | 0.795767002320066 | No |
| *dfrA16* | 1,01535494 | 0.0153549358213231 | 0.394351855702621 | No |
| *dfrA17* | 0,99246385 | -0.00753614816675217 | 0.675851191837805 | No |
| *dfrA21* | 1,07490992 | 0.0749099184370004 | 3.66651887357442e-05 | Yes |
| *dfrA22* | 1,03212833 | 0.0321283284915967 | 0.0750105142463746 | No |
| *dfrA25* | 1,05814018 | 0.0581401758049288 | 0.00131629611988486 | Yes |
| *dfrA27* | 1,02656295 | 0.0265629546000959 | 0.140859365699586 | No |
| *dfrA29* | 1,02421651 | 0.0242165074038749 | 0.179363510517951 | No |
| *dfrA30* | 0,99651632 | -0.00348367867611084 | 0.846726162252624 | No |
| *dfrA31* | 0,85184263 | -0.148157369069202 | 1.18739642917594e-15 | Yes |
| *dfrA34* | 1,04931161 | 0.0493116066120347 | 0.00637531062544367 | Yes |
| *dfrA35* | 0,96199716 | -0.0380028398372901 | 0.0353045310506941 | Yes |
| *dfrA5* | 0,96908404 | -0.0309159596795574 | 0.0866424763668795 | No |
| *dfrA6* | 1,02108131 | 0.021081313133214 | 0.242366807452931 | No |
| *dfrA7* | 0,90058469 | -0.099415313977862 | 5.05809320804145e-08 | Yes |
| *dfrB1* | 0,99730654 | -0.00269346090930935 | 0.881197183751999 | No |
| *dfrB2* | 0,88879334 | -0.111206655118373 | 1.22192789591245e-09 | Yes |
| *dfrB3* | 0,97810761 | -0.0218923928970861 | 0.2247410229117 | No |
| *dfrB4* | 0,86800991 | -0.131990086629749 | 7.50355891909296e-13 | Yes |
| *dfrB5* | 0,96913629 | -0.0308637119721874 | 0.0871748748572427 | No |
| *dfrB6* | 1,02558654 | 0.0255865444758025 | 0.156027615379975 | No |
| *dfrB7* | 1,02445992 | 0.0244599222975162 | 0.175037579653919 | No |
| *dfrB8* | 0,95412961 | -0.0458703919764445 | 0.0111327288046359 | Yes |
| *dfrB9* | 0,95678711 | -0.0432128897459212 | 0.0167502016601731 | Yes |
| *ereA2* | 1,38762587 | 0.38762587456872 | 7.67008572196664e-77 | Yes |
| *ereA3* | 1,11887879 | 0.118878785784974 | 8.99701798216223e-11 | Yes |
| *fosC2* | 0,96886364 | -0.0311363613446677 | 0.0844255069637775 | No |
| *fosE* | 0,98465037 | -0.015349630827938 | 0.394515142043139 | No |
| *fosF* | 0,88757918 | -0.112420816510102 | 8.16474150365988e-10 | Yes |
| *fosG* | 0,94302882 | -0.0569711796458333 | 0.00164200427103464 | Yes |
| *fosH* | 0,96801631 | -0.0319836909442028 | 0.0763272561128757 | No |
| *fosI* | 1,07381139 | 0.0738113869547571 | 4.74511907089172e-05 | Yes |
| *fosK* | 0,88472193 | -0.115278069497636 | 3.11626055511195e-10 | Yes |
| *fosL* | 1,02924211 | 0.0292421127113081 | 0.105056108554606 | No |
| *fosM* | 1,03615769 | 0.036157694727816 | 0.045177732486547 | Yes |
| *fosN* | 0,906155 | -0.0938450031905921 | 2.59567026405454e-07 | Yes |
| *qacE* | 0,86503282 | -0.134967176259973 | 2.39241089445013e-13 | Yes |
| *qacF* | 1,00211936 | 0.00211936138174216 | 0.906386853031281 | No |
| *qacG* | 0,89036283 | -0.109637166146538 | 2.04661042067611e-09 | Yes |
| *qacL* | 1,02038906 | 0.0203890622720161 | 0.258163423701295 | No |
| *qacM* | 0,87888029 | -0.121119707127228 | 4.0870501311157e-11 | Yes |
| *sat2* | 0,9463741 | -0.0536259000759962 | 0.00302839738685803 | Yes |
| *smr2* | 0,96642316 | -0.0335768426742675 | 0.0628222722142912 | No |
| *smr3* | 0,96663504 | -0.0333649572469046 | 0.0644953733829977 | No |

**Supplementary Table 2. Fitness effect of ARC families.** Using a linear regression model we estimate the coefficient of variation in the fitness of the host due to presence of each family of ARCs and its significance (Pr(>|t|) = p-value associated with the t-value). *qacEDsul1* was excluded of the analysis.

| Family | Estimate Coefficient | Std. Error | t value | Pr(>\|t\|) | Signif. |
|---|---|---|---|---|---|
| aminoglycoside | 0.0031229 | 0.0053266 | 0.586 | 0.557873 | ns |
| anti folates | -0.0146423 | 0.0075330 | -1.944 | 0.052312 | ns |
| beta lactam | -0.1037479 | 0.0104613 | -9.917 | < 2.00e-16 | *** |
| chloramphenicol | -0.0627628 | 0.0175051 | -3.585 | 0.000359 | *** |
| fosfomycin | -0.0317774 | 0.0123780 | -2.567 | 0.010451 | * |
| macrolides | 0.2532523 | 0.0276780 | 9.150 | < 2.00e-16 | *** |
| QACs | -0.0585939 | 0.0147945 | -3.961 | 8.22e-05 | *** |
| rifampicin | 0.0007564 | 0.0225990 | 0.033 | 0.973307 | ns |

**Supplementary Table 3. Fitness of ARCs in anaerobiosis.** Using a linear regression model we estimate the coefficient of variation in the fitness of the host due to presence of the ARC and its significance (p-value). *qacEDsul1* was excluded of the analysis.

| ARC | Mean Fitness | Estimate Coefficient | p.value | Signif. |
|---|---|---|---|---|
| *aacA16* | 0.88818546 | -0.111814542368227 | 1.43020812086819e-39 | Yes |
| *aacA2* | 1.02540391 | 0.0254039137648204 | 0.00139501562406131 | Yes |
| *aacA27* | 0.97367978 | -0.0263202183702418 | 0.000933071004145589 | Yes |
| *aacA28* | 0.91546436 | -0.0845356444439471 | 1.36268006952517e-24 | Yes |
| *aacA29* | 0.99264448 | -0.00735552268996281 | 0.353027324617999 | No |
| *aacA3* | 1.0102755 | 0.0102755008510779 | 0.194632115328611 | No |
| *aacA30* | 0.91206294 | -0.0879370633637789 | 2.56428018813697e-26 | Yes |
| *aacA31* | 0.99350312 | -0.00649687716293009 | 0.411999777427032 | No |
| *aacA34* | 0.97942948 | -0.0205705174493173 | 0.00956227794842338 | Yes |
| *aacA35* | 0.98541257 | -0.0145874270084295 | 0.0657663808371128 | No |
| *aacA37* | 1.00988888 | 0.00988887855814143 | 0.21193642058439 | No |
| *aacA38* | 1.00499908 | 0.00499907700410183 | 0.527833421697994 | No |
| *aacA4* | 1.00579365 | 0.00579365317221418 | 0.464399666266899 | No |
| *aacA42* | 0.92407012 | -0.0759298813532968 | 1.94275257108815e-20 | Yes |
| *aacA43* | 0.88881306 | -0.111186936391225 | 3.35898473895961e-39 | Yes |
| *aacA45* | 1.00367062 | 0.00367062336197947 | 0.642945390494406 | No |
| *aacA47* | 0.97789862 | -0.0221013781619788 | 0.00538637176210709 | Yes |
| *aacA48* | 1.00851725 | 0.00851724719259447 | 0.282244409204175 | No |
| *aacA50* | 0.99377617 | -0.00622383353973016 | 0.431915057815362 | No |
| *aacA51* | 0.99283878 | -0.0071612164403353 | 0.365881057727054 | No |
| *aacA52* | 0.97958646 | -0.0204135449719018 | 0.0101230121031668 | Yes |
| *aacA54* | 1.01539589 | 0.0153958927977411 | 0.0521742497138952 | No |
| *aacA59* | 1.01042849 | 0.0104284870748967 | 0.188079671262742 | No |
| *aacA61* | 0.9669787 | -0.0330212952109053 | 3.43656575362306e-05 | Yes |
| *aacA64* | 0.92363725 | -0.0763627489462424 | 1.22192685242272e-20 | Yes |
| *aacA7* | 0.94691567 | -0.0530843346220288 | 4.40919580586168e-11 | Yes |
| *aacA8* | 0.91358511 | -0.0864148890311938 | 1.53716301695987e-25 | Yes |
| *aacAX* | 0.95383848 | -0.0461615247038747 | 8.71438220872600e-09 | Yes |
| *aacC1* | 0.99433843 | -0.00566156918703544 | 0.47464072166871 | No |
| *aacC11* | 0.98852831 | -0.0114716908604802 | 0.147688338915958 | No |
| *aacC13* | 0.91705522 | -0.0829447827999136 | 8.42551159179977e-24 | Yes |
| *aacC2* | 0.99418269 | -0.00581731324317112 | 0.46257830171021 | No |
| *aacC3* | 0.97508096 | -0.0249190435115458 | 0.00171757834495151 | Yes |
| *aacC4* | 1.00013612 | 0.000136119451329872 | 0.986282813495671 | No |
| *aacC5* | 0.95455059 | -0.0454494064384719 | 1.44894990239103e-08 | Yes |
| *aacC6* | 0.99992852 | -7.14817372971172e-05 | 0.992796302216256 | No |
| *aadA1* | 1.01025506 | 0.0102550571451573 | 0.195520287992696 | No |
| *aadA10* | 1.0079255 | 0.00792550404419471 | 0.316998018507628 | No |
| *aadA11* | 0.99858087 | -0.00141912568094984 | 0.857747110984892 | No |
| *aadA13* | 0.96988904 | -0.0301109576768169 | 0.000155799958532185 | Yes |
| *aadA16* | 0.99207137 | -0.00792862602511145 | 0.316807575560758 | No |
| *aadA2* | 0.98929202 | -0.0107079761651898 | 0.176532819633306 | No |
| *aadA24* | 0.95462003 | -0.0453799697677697 | 1.52204999983604e-08 | Yes |
| *aadA28* | 0.97472002 | -0.025279979216736 | 0.00147165874371937 | Yes |
| *aadA29* | 1.00612306 | 0.00612305531144821 | 0.439404585931945 | No |
| *aadA34* | 1.0126278 | 0.0126278024319779 | 0.111077699958758 | No |
| *aadA4* | 1.00889795 | 0.00889794736160617 | 0.261307170799104 | No |
| *aadA5* | 1.0226573 | 0.0226573022571856 | 0.00433731925123746 | Yes |
| *aadA6* | 1.0288305 | 0.0288304989841986 | 0.00029174098002905 | Yes |
| *aadA7* | 1.0324082 | 0.0324082041596407 | 4.77198843051302e-05 | Yes |
| *aadB* | 0.958875 | -0.0411250016847504 | 2.74789605321441e-07 | Yes |
| *aphA15* | 1.00987136 | 0.00987136029531336 | 0.212746081789463 | No |
| *aphA16* | 0.74709522 | -0.252904775593508 | 6.18462127744657e-134 | Yes |
| *arr5* | 0.99600853 | -0.00399147271908638 | 0.614192018994877 | No |
| *arr6* | 0.99834312 | -0.00165687837984243 | 0.834236588669039 | No |
| *arr7* | 1.05302307 | 0.0530230676470966 | 4.6328169904731e-11 | Yes |
| *bla*$_{BEL-1}$ | 0.99043569 | -0.00956431194628001 | 0.227302350855701 | No |
| *bla*$_{GES-1}$ | 0.97212597 | -0.0278740273625403 | 0.000459136743014409 | Yes |
| *bla*$_{IMP-2}$ | 0.9328877 | -0.0671122990902588 | 1.60046109984009e-16 | Yes |
| *bla*$_{IMP-31}$ | 0.81904227 | -0.180957727111518 | 1.02444432157664e-84 | Yes |
| *bla*$_{OXA-1}$ | 0.89257766 | -0.107422337804334 | 5.34004917572467e-37 | Yes |
| *bla*$_{OXA-10}$ | 0.98954282 | -0.0104571789226861 | 0.186869175345832 | No |
| *bla*$_{OXA-118}$ | 0.96666774 | -0.0333322550856628 | 2.90360973513416e-05 | Yes |

| ARC | Mean Fitness | Estimate Coefficient | p.value | Signif. |
|---|---|---|---|---|
| *bla*$_{OXA-129}$ | 0.86140355 | -0.138596447890502 | 3.03763093485754e-56 | Yes |
| *bla*$_{OXA-198}$ | 1.00065872 | 0.000658717271177778 | 0.933692395623497 | No |
| *bla*$_{OXA-2}$ | 0.84485085 | -0.155149151366856 | 3.61838218923809e-67 | Yes |
| *bla*$_{OXA-46}$ | 0.61562764 | -0.384372355315825 | 3.64345003177508e-215 | Yes |
| *bla*$_{OXA-5}$ | 0.91446074 | -0.0855392591612392 | 4.26604962038884e-25 | Yes |
| *bla*$_{OXA-9}$ | 0.96595684 | -0.0340431649841392 | 1.96525403165753e-05 | Yes |
| *bla*$_{VIM-1}$ | 0.92445646 | -0.0755435373957293 | 2.93390400800848e-20 | Yes |
| *bla*$_{VIM-2}$ | 0.90121478 | -0.0987852223377399 | 4.15777061650199e-32 | Yes |
| *bla*$_{VIM-7}$ | 0.98132152 | -0.0186784792847363 | 0.0185712690512566 | Yes |
| *catB10* | 0.98875539 | -0.0112446114330192 | 0.155860064470431 | No |
| *catB2* | 0.84484836 | -0.155151636979202 | 3.60453547867108e-67 | Yes |
| *catB3* | 0.97862596 | -0.0213740369481794 | 0.00710422764616624 | Yes |
| *catB5* | 0.94079008 | -0.0592099165777994 | 2.47861706524161e-13 | Yes |
| *catB6* | 0.95846483 | -0.0415351734228553 | 2.10120507776678e-07 | Yes |
| *dfrA1* | 0.9314048 | -0.0685951986049421 | 3.72562104453289e-17 | Yes |
| *dfrA12* | 0.93761314 | -0.0623868600863455 | 1.41133638449275e-14 | Yes |
| *dfrA14* | 0.98123325 | -0.0187667486703357 | 0.0180250965326888 | Yes |
| *dfrA15* | 0.96043011 | -0.0395698857658761 | 7.4441402029789e-07 | Yes |
| *dfrA16* | 0.95745544 | -0.0425445586600429 | 1.07521956640133e-07 | Yes |
| *dfrA17* | 0.94573028 | -0.0542697159671747 | 1.67737990540927e-11 | Yes |
| *dfrA21* | 1.01100467 | 0.0110046690327404 | 0.164863654129723 | No |
| *dfrA22* | 1.01288435 | 0.0128843512323483 | 0.104019490833296 | No |
| *dfrA25* | 0.98146442 | -0.0185355827627596 | 0.0194862437225526 | Yes |
| *dfrA27* | 0.94582126 | -0.0541787353321734 | 1.80766088756234e-11 | Yes |
| *dfrA29* | 0.99677894 | -0.00322106032641007 | 0.684144145522116 | No |
| *dfrA30* | 0.9835912 | -0.0164087975123063 | 0.0385443691743575 | Yes |
| *dfrA31* | 0.91210711 | -0.0878928862562116 | 2.70186653500056e-26 | Yes |
| *dfrA34* | 0.99342178 | -0.00657822229842886 | 0.406173848704657 | No |
| *dfrA35* | 0.96494698 | -0.0350530240017476 | 1.11520947663671e-05 | Yes |
| *dfrA5* | 0.96560283 | -0.0343971679164617 | 1.61385278720123e-05 | Yes |
| *dfrA6* | 0.91011675 | -0.0898832519316878 | 2.5213410312369e-27 | Yes |
| *dfrA7* | 0.85895539 | -0.141044614217571 | 7.76883546624718e-58 | Yes |
| *dfrB1* | 0.9753185 | -0.0246814957444245 | 0.00189953527242418 | Yes |
| *dfrB2* | 0.99968939 | -0.000310611594049585 | 0.968705191435775 | No |
| *dfrB3* | 0.95831425 | -0.0416857459717304 | 1.90301488965945e-07 | Yes |
| *dfrB4* | 0.8953804 | -0.104619601069559 | 2.18735877718062e-35 | Yes |
| *dfrB5* | 0.99950566 | -0.000494338187256156 | 0.950213949249907 | No |
| *dfrB6* | 0.97820464 | -0.0217953638441115 | 0.00605719980767237 | Yes |
| *dfrB7* | 0.99670608 | -0.00329391880620363 | 0.677398822889152 | No |
| *dfrB8* | 0.95822738 | -0.0417726173918437 | 1.79704151672149e-07 | Yes |
| *dfrB9* | 0.9665808 | -0.0334191985370282 | 2.76930377721066e-05 | Yes |
| *ereA2* | 0.86579955 | -0.13420045279783 | 2.07627886402619e-53 | Yes |
| *ereA3* | 0.92107491 | -0.0789250933588724 | 7.55519758133371e-22 | Yes |
| *fosC2* | 0.94668006 | -0.0533199374306314 | 3.64371738998041e-11 | Yes |
| *fosE* | 0.97883912 | -0.0211608842201547 | 0.00769368104971156 | Yes |
| *fosF* | 0.97377201 | -0.0262279883694019 | 0.000972140266458111 | Yes |
| *fosG* | 0.97618689 | -0.023813109051419 | 0.00272621418537525 | Yes |
| *fosH* | 0.96267303 | -0.0373269744803273 | 2.95663833090814e-06 | Yes |
| *fosI* | 0.99130168 | -0.00869831973967378 | 0.272147884466648 | No |
| *fosK* | 0.94557248 | -0.0544275231752081 | 1.47291787769473e-11 | Yes |
| *fosL* | 1.00401108 | 0.00401107651541484 | 0.612453727869469 | No |
| *fosM* | 0.99647252 | -0.00352747902760179 | 0.655951981131507 | No |
| *fosN* | 0.97101492 | -0.0289850788667577 | 0.000270795770557815 | Yes |
| *qacE* | 1.0070819 | 0.00708190093170108 | 0.371211025323446 | No |
| *qacF* | 0.96823538 | -0.0317646207384475 | 6.69739338355342e-05 | Yes |
| *qacG* | 0.97803899 | -0.0219610072661738 | 0.00568526174679478 | Yes |
| *qacH* | 0.97951269 | -0.0204873100288299 | 0.00985595402413285 | Yes |
| *qacK* | 0.93506895 | -0.064931049094233 | 1.30612904285598e-15 | Yes |
| *qacL* | 0.99450888 | -0.00549111581826872 | 0.488038452064334 | No |
| *qacM* | 0.98680104 | -0.0131989592646219 | 0.095856719311087 | No |
| *sat2* | 1.00151807 | 0.00151807116238617 | 0.847946111947095 | No |
| *smr2* | 0.96688736 | -0.0331126423638163 | 3.27104401802226e-05 | Yes |
| *smr3* | 0.95595056 | -0.0440494350231135 | 3.86063287712273e-08 | Yes |

**Supplementary Table 4. Fitness effect of ARC families in anaerobiosis.** Using a linear regression model we estimate the coefficient of variation in the fitness of the host due to presence of each family of ARCs and its significance (Pr(>|t|) = p-value associated with the t-value). *qacEDsul1* was excluded of the analysis.

| Family | Estimate Coefficient | Std. Error | t value | Pr(>\|t\|) | Signif. |
|---|---|---|---|---|---|
| aminoglycoside | -0.023208 | 0.002982 | -7.782 | 2.38e-14 | *** |
| anti folates | -0.037834 | 0.004218 | -8.970 | < 2.00e-16 | *** |
| beta lactam | -0.089173 | 0.005479 | -16.276 | < 2.00e-16 | *** |
| chloramphenicol | -0.057703 | 0.009801 | -5.888 | 5.92e-09 | *** |
| fosfomycin | -0.025348 | 0.006930 | -3.658 | 0.000273 | *** |
| macrolides | -0.106563 | 0.015497 | -6.877 | 1.30e-11 | *** |
| QACs | -0.025324 | 0.007305 | -3.467 | 0.000557 | *** |
| rifampicin | 0.015792 | 0.012653 | 1.248 | 0.212402 | ns |

**Supplementary Table 5. Effect of oxygen in ARC families fitness effect.** We estimate for each family of ARCs the variation of the fitness effect depending on oxygen availability. Using a linear regression model we determine the significance (Pr(>|t|) = p-value associated with the t-value). *qacEDsul1* was excluded of the analysis together with ARCs that were not able to be measure in both conditions.

| Family | Estimate Coefficient | Std. Error | t value | Pr(>\|t\|) | Signif. |
|---|---|---|---|---|---|
| ARCs | -0.018671 | 0.004475 | -4.173 | 3.19e-05 | *** |
| aminoglycoside | -0.0263312 | 0.0060786 | -4.332 | 1.58e-05 | *** |
| anti folates | -0.0231914 | 0.0085965 | -2.698 | 0.00706 | ** |
| beta lactam | 0.0145748 | 0.0115591 | 1.261 | 0.20755 | ns |
| chloramphenicol | 0.0050597 | 0.0199764 | 0.253 | 0.80009 | ns |
| fosfomycin | 0.0064297 | 0.0141255 | 0.455 | 0.64904 | ns |
| macrolides | -0.3598151 | 0.0315855 | -11.392 | < 2.00e-16 | *** |
| QACs | 0.0332701 | 0.0159176 | 2.090 | 0.03678 | * |
| rifampicin | 0.0150351 | 0.0257894 | 0.583 | 0.55999 | ns |

**Supplementary Table 6. Effect of oxygen in ARCs fitness effect.** We estimate for each family of ARCs the estimate coefficient of variation of the fitness effect depending on oxygen availability (Anaero – aero). Using a linear regression model we determined its significance (p-value). *qacEDsul1* was excluded of the analysis together with ARCs that were not able to be measure in both conditions.

| ARC | Estimate Coefficient | p.value | Signif. |
|---|---|---|---|
| *aacA16* | -0.00284487645307367 | 0.884436116333142 | No |
| *aacA2* | -0.0650741961465879 | 0.000908746715387576 | Yes |
| *aacA27* | -0.120326914552796 | 1.05034580157022e-09 | Yes |
| *aacA28* | 0.0257653760509293 | 0.188198033208357 | No |
| *aacA29* | 0.0146662356978293 | 0.453716677379749 | No |
| *aacA3* | -0.0894305477583754 | 5.36295680362473e-06 | Yes |
| *aacA30* | 0.0698161447091021 | 0.000373832327821233 | Yes |
| *aacA31* | -0.0729138182345187 | 0.000203281790877981 | Yes |
| *aacA34* | -0.093498542143531 | 1.98250069600657e-06 | Yes |
| *aacA35* | 0.0191444102295249 | 0.328111217162386 | No |
| *aacA37* | -0.00313434551344451 | 0.872773026761822 | No |
| *aacA38* | 0.00118403193673469 | 0.951762382713243 | No |
| *aacA4* | -0.065945720312579 | 0.00077497581027666 | Yes |
| *aacA42* | -0.226502629615538 | 1.70783186747207e-29 | Yes |
| *aacA45* | -0.0984839131313188 | 5.54961387398044e-07 | Yes |
| *aacA47* | -0.136207674964186 | 5.48323832318276e-12 | Yes |
| *aacA48* | -0.0244663390230433 | 0.211436563267248 | No |
| *aacA50* | -0.019967505359923 | 0.307750714933316 | No |
| *aacA51* | 0.0446540678446577 | 0.0226642233679374 | Yes |
| *aacA52* | -0.0125510610495304 | 0.52139426029353 | No |
| *aacA54* | 0.0175915147315312 | 0.368850769248485 | No |
| *aacA59* | -0.0913350163785547 | 3.38220237121227e-06 | Yes |
| *aacA61* | -0.100207745105269 | 3.52458399585289e-07 | Yes |
| *aacA64* | 0.0422670417893293 | 0.0309703084305755 | Yes |
| *aacA7* | -0.126936646100377 | 1.26482333732943e-10 | Yes |
| *aacA8* | -0.152526945627471 | 1.36259491088327e-14 | Yes |
| *aacAX* | 0.0861999491721152 | 1.14930139271371e-05 | Yes |
| *aacC1* | 0.00760743619186206 | 0.697525855514186 | No |
| *aacC11* | -0.048683078417771 | 0.012984654999401 | Yes |
| *aacC13* | -0.00537738513342354 | 0.783520787469192 | No |
| *aacC2* | -0.104215731701061 | 1.19374399763984e-07 | Yes |
| *aacC3* | 0.0776421561845268 | 7.67460935533939e-05 | Yes |
| *aacC4* | 0.140329646322939 | 1.2755935376123e-12 | Yes |
| *aacC5* | -0.0123017084405863 | 0.529701977302284 | No |
| *aacC6* | 0.0545302323760349 | 0.00540787617850619 | Yes |
| *aadA1* | 0.0304411096658207 | 0.12006044805106 | No |
| *aadA10* | 0.00541206321788533 | 0.782159885592279 | No |
| *aadA11* | -0.154996254450225 | 5.22098785325491e-15 | Yes |
| *aadA13* | 0.053068007267631 | 0.00678308851817419 | Yes |
| *aadA16* | 0.0182533430576519 | 0.351117085401569 | No |
| *aadA2* | -0.0301342617676741 | 0.1238371792989 | No |
| *aadA24* | 0.0338387981951157 | 0.0840187864551999 | No |
| *aadA28* | 0.0139763663370698 | 0.475228279050906 | No |
| *aadA29* | 0.0232052214000127 | 0.23591839745248 | No |
| *aadA34* | 0.00672506973474582 | 0.731158546797249 | No |
| *aadA4* | -0.022324684567184 | 0.254161539914554 | No |
| *aadA5* | -0.0841926804646524 | 1.82250357638986e-05 | Yes |
| *aadA6* | -0.0221985669818841 | 0.256852844426821 | No |
| *aadA7* | 0.0022115291668971 | 0.91003867862303 | No |
| *aadB* | 0.174313382634825 | 1.81574979818727e-18 | Yes |
| *aphA15* | -0.0333213997980336 | 0.0888600304939032 | No |
| *aphA16* | -0.435812564789748 | 1.06920747549162e-92 | Yes |
| *arr5* | 0.0124635729509989 | 0.52430145061447 | No |
| *arr6* | 0.00124453580468576 | 0.949300690504569 | No |
| *arr7* | 0.0313972697163082 | 0.108868918706286 | No |
| *bla*$_{BEL-1}$ | -0.0890278578633843 | 5.90549157119048e-06 | Yes |
| *bla*$_{GES-1}$ | -0.0499376065867739 | 0.0108329919153738 | Yes |
| *bla*$_{IMP-2}$ | 0.00484684259977234 | 0.804421669253792 | No |
| *bla*$_{IMP-31}$ | 0.0859248596721936 | 1.22496093336582e-05 | Yes |
| *bla*$_{OXA-1}$ | -0.175008269610212 | 1.34318676848518e-18 | Yes |

| ARC | Estimate Coefficient | p.value | Signif. |
|---|---|---|---|
| *bla*$_{OXA-10}$ | -0.107741383469151 | 4.46361370246674e-08 | Yes |
| *bla*$_{OXA-118}$ | 0.233665872595269 | 3.45787981013065e-31 | Yes |
| *bla*$_{OXA-129}$ | 0.0714811760769699 | 0.000270208834769765 | Yes |
| *bla*$_{OXA-198}$ | -0.0391619174361892 | 0.0455842115323843 | Yes |
| *bla*$_{OXA-2}$ | 0.113397067157768 | 8.6664583846207e-09 | Yes |
| *bla*$_{VIM-1}$ | 0.232754491597386 | 5.70933110391544e-31 | Yes |
| *bla*$_{VIM-2}$ | 0.187788115463481 | 4.38140119350971e-21 | Yes |
| *bla*$_{VIM-7}$ | 0.0709952366454826 | 0.000297262878656778 | Yes |
| *catB10* | 0.00800426186044921 | 0.682587455173101 | No |
| *catB2* | -0.0660070596927645 | 0.000766286774245292 | Yes |
| *catB3* | 0.0648446191735972 | 0.000947390239727463 | Yes |
| *catB5* | 0.000388205801512052 | 0.984175824776274 | No |
| *catB6* | 0.018068354807752 | 0.356018505161623 | No |
| *dfrA1* | -0.0945279912399439 | 1.53155413610866e-06 | Yes |
| *dfrA12* | -0.0169781863419467 | 0.385773753284501 | No |
| *dfrA14* | -0.0110532943497777 | 0.572283561323265 | No |
| *dfrA15* | -0.0349051079276331 | 0.0747154255174276 | No |
| *dfrA16* | -0.0578994944813658 | 0.0031473931320919 | Yes |
| *dfrA17* | -0.0467335678004225 | 0.0170817548481403 | Yes |
| *dfrA21* | -0.06390524940426 | 0.00112192779682438 | Yes |
| *dfrA22* | -0.0192439772592483 | 0.325602779554605 | No |
| *dfrA25* | -0.0766757585676883 | 9.40599849605494e-05 | Yes |
| *dfrA27* | -0.0807416899322692 | 3.93694085969441e-05 | Yes |
| *dfrA29* | -0.027437567730285 | 0.161134037328204 | No |
| *dfrA30* | -0.0129251188361955 | 0.509058827081645 | No |
| *dfrA31* | 0.0602644828129905 | 0.00211824760579432 | Yes |
| *dfrA34* | -0.0558898289104636 | 0.00436082145501692 | Yes |
| *dfrA35* | 0.00294981583554247 | 0.880204935684559 | No |
| *dfrA5* | -0.00348120823690439 | 0.858834066417399 | No |
| *dfrA6* | -0.110964565064902 | 1.77010802435699e-08 | Yes |
| *dfrA7* | -0.0416293002397085 | 0.0335898543373227 | Yes |
| *dfrB1* | -0.0219880348351152 | 0.261389529624734 | No |
| *dfrB2* | 0.110896043524323 | 1.80571598083107e-08 | Yes |
| *dfrB3* | -0.0197933530746443 | 0.311987259210718 | No |
| *dfrB4* | 0.02737048556019 | 0.162159787233853 | No |
| *dfrB5* | 0.0303693737849312 | 0.120935194621041 | No |
| *dfrB6* | -0.0473819083199133 | 0.0156081134607213 | Yes |
| *dfrB7* | -0.0277538411037196 | 0.156363929834982 | No |
| *dfrB8* | 0.00409777458460094 | 0.834167959688196 | No |
| *dfrB9* | 0.00979369120889303 | 0.616830208156775 | No |
| *ereA2* | -0.52182632736655 | 3.34029321251897e-124 | Yes |
| *ereA3* | -0.197803879143846 | 3.89257920888751e-23 | Yes |
| *fosC2* | -0.0221835760859636 | 0.257174057056693 | No |
| *fosE* | -0.00581125339221667 | 0.766543168012975 | No |
| *fosF* | 0.0861928281406999 | 1.15120234940175e-05 | Yes |
| *fosG* | 0.0331580705944144 | 0.0904341953867629 | No |
| *fosH* | -0.00534328353612449 | 0.784859712701249 | No |
| *fosI* | -0.0825097066944309 | 2.66277641405159e-05 | Yes |
| *fosK* | 0.0608505463224285 | 0.00191632717639773 | Yes |
| *fosL* | -0.0252310361958931 | 0.197517575993515 | No |
| *fosM* | -0.0396851737554178 | 0.0427758911418548 | Yes |
| *fosN* | 0.0648599243238344 | 0.000944767321313457 | Yes |
| *qacE* | 0.142049077191674 | 6.86424941104712e-13 | Yes |
| *qacF* | -0.0338839821201895 | 0.0836063556302897 | No |
| *qacG* | 0.0876761588803638 | 8.13788873233178e-06 | Yes |
| *qacL* | -0.0258801780902848 | 0.186238838699504 | No |
| *qacM* | 0.107920747862606 | 4.24240449436048e-08 | Yes |
| *sat2* | 0.0551439712383824 | 0.00490990388112978 | Yes |
| *smr2* | 0.000464200310451162 | 0.981078644948404 | No |
| *smr3* | -0.0106844777762089 | 0.585167844313377 | No |

**Supplementary Table 7. Strains and plasmids used in this study.** All strains used in this work were built in *E. coli* backgrounds otherwise stated. Strains corresponding to the OMM[12] consortium are numbered following DSM (German collection of microorganisms and cell cultures) classification.

| Strain | Genetic background | Plasmid | ARC cloned (family) | Source |
|---|---|---|---|---|
| | | **General purpose strains** | | |
| A072 | MG1655 | - | - | Lab collection |
| A093 | DH5α | - | - | " |
| A249 | MG1655 | pMBA empty vector | - | Hipólito et al. 2023 |
| C381 | DH5α | pMBA empty vector | - | " |
| | | **pMBA collection used for flow cytometry competition assays** | | |
| A223 | MG1655 | pMBA$_{dfrA1}$ | *dfrA1*(dfr) | " |
| A224 | MG1655 | pMBA$_{dfrA5}$ | *dfrA5* (dfr) | " |
| A225 | MG1655 | pMBA$_{dfrA6}$ | *dfrA6* (dfr) | " |
| A226 | MG1655 | pMBA$_{dfrA7}$ | *dfrA7* (dfr) | " |
| A241 | MG1655 | pMBA$_{dfrA12}$ | *dfrA12* (dfr) | " |
| A227 | MG1655 | pMBA$_{dfrA14}$ | *dfrA14* (dfr) | " |
| A228 | MG1655 | pMBA$_{dfrA15}$ | *dfrA15* (dfr) | " |
| A229 | MG1655 | pMBA$_{dfrA16}$ | *dfrA16* (dfr) | " |
| A230 | MG1655 | pMBA$_{dfrA17}$ | *dfrA17* (dfr) | " |
| A231 | MG1655 | pMBA$_{dfrA21}$ | *dfrA21*(dfr) | " |
| A500 | MG1655 | pMBA$_{dfrA22}$ | *dfrA22* (dfr) | " |
| A244 | MG1655 | pMBA$_{dfrA22.2}$ | *dfrA22.2* (dfr) | " |
| A232 | MG1655 | pMBA$_{dfrA25}$ | *dfrA25* (dfr) | " |
| A233 | MG1655 | pMBA$_{dfrA27}$ | *dfrA27*(dfr) | " |
| A613 | MG1655 | pMBA$_{dfrA29}$ | *dfrA29* (dfr) | " |
| A248 | MG1655 | pMBA$_{dfrA30}$ | *dfrA30* (dfr) | " |
| A234 | MG1655 | pMBA$_{dfrA31}$ | *dfrA31* (dfr) | " |
| A235 | MG1655 | pMBA$_{dfrA34}$ | *dfrA34* (dfr) | " |
| A242 | MG1655 | pMBA$_{dfrA35}$ | *dfrA35* (dfr) | " |
| A236 | MG1655 | pMBA$_{dfrB1}$ | *dfrB1* (dfr) | " |
| A245 | MG1655 | pMBA$_{dfrB2}$ | *dfrB2* (dfr) | " |
| A246 | MG1655 | pMBA$_{dfrB3}$ | *dfrB3* (dfr) | " |
| A237 | MG1655 | pMBA$_{dfrB4}$ | *dfrB4* (dfr) | " |
| A238 | MG1655 | pMBA$_{dfrB5}$ | *dfrB5* (dfr) | " |
| A239 | MG1655 | pMBA$_{dfrB6}$ | *dfrB6* (dfr) | " |
| A240 | MG1655 | pMBA$_{dfrB7}$ | *dfrB7* (dfr) | " |
| A247 | MG1655 | pMBA$_{dfrB8}$ | *dfrB8* (dfr) | " |
| A243 | MG1655 | pMBA$_{dfrB9}$ | *dfrB9* (dfr) | " |
| A390 | MG1655 | pMBA$_{blaBEL-1}$ | *bla*$_{BEL-1}$ (bla) | " |
| A380 | MG1655 | pMBA$_{blaGES-1}$ | *bla*$_{GES-1}$ (bla) | " |
| A387 | MG1655 | pMBA$_{blaIMP-2}$ | *bla*$_{IMP-2}$ (bla) | " |
| A383 | MG1655 | pMBA$_{blaIMP-31}$ | *bla*$_{IMP-31}$ (bla) | " |
| A441 | MG1655 | pMBA$_{blaOXA-1}$ | *bla*$_{OXA-1}$ (bla) | " |
| A373 | MG1655 | pMBA$_{blaOXA-2}$ | *bla*$_{OXA-2}$ (bla) | " |
| A381 | MG1655 | pMBA$_{blaOXA-5}$ | *bla*$_{OXA-5}$ (bla) | " |
| A384 | MG1655 | pMBA $_{blaOXA-9}$ | *bla*$_{OXA-9}$ (bla) | " |
| A374 | MG1655 | pMBA$_{blaOXA-10}$ | *bla*$_{OXA-10}$ (bla) | " |
| A385 | MG1655 | pMBA$_{blaOXA-20}$ | *bla*$_{OXA-20}$ (bla) | " |
| A382 | MG1655 | pMBA$_{blaOXA-21}$ | *bla*$_{OXA-21}$ (bla) | " |
| A395 | MG1655 | pMBA$_{blaOXA-46}$ | *bla*$_{OXA-46}$ (bla) | " |
| A375 | MG1655 | pMBA$_{blaOXA-118}$ | *bla*$_{OXA-118}$ (bla) | " |
| A391 | MG1655 | pMBA$_{blaOXA-129}$ | *bla*$_{OXA-129}$ (bla) | " |
| A376 | MG1655 | pMBA$_{blaOXA-198}$ | *bla*$_{OXA-198}$ (bla) | " |
| A803 | MG1655 | pMBA$_{blaPBL-1}$ | *bla*$_{PBL-1}$ (bla) | " |
| A388 | MG1655 | pMBA$_{blaVIM-1}$ | *bla*$_{VIM-1}$ (bla) | " |
| A396 | MG1655 | pMBA$_{blaVIM-2}$ | *bla*$_{VIM-2}$ (bla) | " |
| A389 | MG1655 | pMBA$_{blaVIM-7}$ | *bla*$_{VIM-7}$ (bla) | " |
| A327 | MG1655 | pMBA$_{aacA2}$ | *aacA2* (aa) | " |
| A266 | MG1655 | pMBA$_{aacA3}$ | *aacA3* (aa) | " |
| A328 | MG1655 | pMBA$_{aacA4}$ | *aacA4* (aa) | " |
| A267 | MG1655 | pMBA$_{aacA7}$ | *aacA7* (aa) | " |

| | | | | |
|---|---|---|---|---|
| A268 | MG1655 | pMBA$_{aacA8}$ | *aacA8* (aa) | “ |
| A263 | MG1655 | pMBA$_{aacA16}$ | *aacA16* (aa) | “ |
| A543 | MG1655 | pMBA$_{aacA17}$ | *aacA17* (aa) | “ |
| A264 | MG1655 | pMBA$_{aacA27}$ | *aacA27* (aa) | “ |
| A256 | MG1655 | pMBA$_{aacA28}$ | *aacA28* (aa) | “ |
| A257 | MG1655 | pMBA$_{aacA29}$ | *aacA29* (aa) | “ |
| A270 | MG1655 | pMBA$_{aacA30}$ | *aacA30* (aa) | “ |
| B657 | MG1655 | pMBA$_{aacA31}$ | *aacA31* (aa) | “ |
| A272 | MG1655 | pMBA$_{aacA34}$ | *aacA34* (aa) | “ |
| A329 | MG1655 | pMBA$_{aacA35}$ | *aacA35* (aa) | “ |
| A258 | MG1655 | pMBA$_{aacA37}$ | *aacA37* (aa) | “ |
| A603 | MG1655 | pMBA$_{aacA38}$ | *aacA38* (aa) | “ |
| A273 | MG1655 | pMBA$_{aacA42}$ | *aacA42* (aa) | “ |
| C117 | MG1655 | pMBA$_{aacA43}$ | *aacA43* (aa) | “ |
| A265 | MG1655 | pMBA$_{aacA45}$ | *aacA45* (aa) | “ |
| A274 | MG1655 | pMBA$_{aacA47}$ | *aacA47* (aa) | “ |
| A259 | MG1655 | pMBA$_{aacA48}$ | *aacA48* (aa) | “ |
| B656 | MG1655 | pMBA$_{aacA49}$ | *aacA49* (aa) | “ |
| A260 | MG1655 | pMBA$_{aacA50}$ | *aacA50* (aa) | “ |
| A261 | MG1655 | pMBA$_{aacA51}$ | *aacA51* (aa) | “ |
| A326 | MG1655 | pMBA$_{aacA52}$ | *aacA52* (aa) | “ |
| B655 | MG1655 | pMBA$_{aacA54}$ | *aacA54* (aa) | “ |
| A262 | MG1655 | pMBA$_{aacA56}$ | *aacA56* (aa) | “ |
| A303 | MG1655 | pMBA$_{aacA59}$ | *aacA59* (aa) | “ |
| A302 | MG1655 | pMBA$_{aacA61}$ | *aacA61* (aa) | “ |
| A566 | MG1655 | pMBA$_{aacA64}$ | *aacA64* (aa) | “ |
| A331 | MG1655 | pMBA$_{aacAX}$ | *aacAX* (aa) | “ |
| A308 | MG1655 | pMBA$_{aacC1}$ | *aacC1* (aa) | “ |
| A304 | MG1655 | pMBA$_{aacC2}$ | *aacC2* (aa) | “ |
| B625 | MG1655 | pMBA$_{aacC3}$ | *aacC3* (aa) | “ |
| A320 | MG1655 | pMBA$_{aacC4}$ | *aacC4* (aa) | “ |
| A305 | MG1655 | pMBA$_{aacC5}$ | *aacC5* (aa) | “ |
| A306 | MG1655 | pMBA$_{aacC6}$ | *aacC6* (aa) | “ |
| A332 | MG1655 | pMBA$_{aacC11}$ | *aacC11* (aa) | “ |
| A309 | MG1655 | pMBA$_{aacC13}$ | *aacC13* (aa) | “ |
| A311 | MG1655 | pMBA$_{aadA1}$ | *aadA1* (aa) | “ |
| A333 | MG1655 | pMBA$_{aadA2}$ | *aadA2* (aa) | “ |
| A319 | MG1655 | pMBA$_{aadA4}$ | *aadA4* (aa) | “ |
| A313 | MG1655 | pMBA$_{aadA5}$ | *aadA5* (aa) | “ |
| A321 | MG1655 | pMBA$_{aadA6}$ | *aadA6* (aa) | “ |
| A397 | MG1655 | pMBA$_{aadA7}$ | *aadA7* (aa) | “ |
| A315 | MG1655 | pMBA$_{aadA10}$ | *aadA10* (aa) | “ |
| A312 | MG1655 | pMBA$_{aadA11}$ | *aadA11* (aa) | “ |
| A322 | MG1655 | pMBA$_{aadA13}$ | *aadA13* (aa) | “ |
| A316 | MG1655 | pMBA$_{aadA16}$ | *aadA16* (aa) | “ |
| A318 | MG1655 | pMBA$_{aadA24}$ | *aadA24* (aa) | “ |
| A604 | MG1655 | pMBA$_{aadA28}$ | *aadA28* (aa) | “ |
| A314 | MG1655 | pMBA$_{aadA29}$ | *aadA29* (aa) | “ |
| A398 | MG1655 | pMBA$_{aadA34}$ | *aadA34* (aa) | “ |
| A292 | MG1655 | pMBA$_{aadB}$ | *aadB* (aa) | “ |
| A422 | MG1655 | pMBA$_{aphA15}$ | *aphA15* (aa) | “ |
| C116 | MG1655 | pMBA$_{aphA16}$ | *aphA16* (aa) | “ |
| A293 | MG1655 | pMBA$_{sat2}$ | *sat2* (aa) | “ |
| A368 | MG1655 | pMBA$_{arr2}$ | *arr2* (mix) | “ |
| A334 | MG1655 | pMBA$_{arr5}$ | *arr5* (mix) | “ |
| A363 | MG1655 | pMBA$_{arr6}$ | *arr6* (mix) | “ |
| A440 | MG1655 | pMBA$_{arr7}$ | *arr7* (mix) | “ |
| A364 | MG1655 | pMBA$_{arr8b}$ | *arr8b* (mix) | “ |
| A335 | MG1655 | pMBA$_{catB2}$ | *catB2* (mix) | “ |
| A336 | MG1655 | pMBA$_{catB3}$ | *catB3* (mix) | “ |
| A365 | MG1655 | pMBA$_{catB5}$ | *catB5* (mix) | “ |
| A625 | MG1655 | pMBA$_{catB6}$ | *catB6* (mix) | “ |
| A393 | MG1655 | pMBA$_{catB10}$ | *catB10* (mix) | “ |
| A338 | MG1655 | pMBA$_{ereA2}$ | *ereA2* (mix) | “ |
| A423 | MG1655 | pMBA$_{ereA3}$ | *ereA3* (mix) | “ |
| A657 | MG1655 | pMBA$_{fosC2}$ | *fosC2* (mix) | “ |

| | | | | |
|---|---|---|---|---|
| A339 | MG1655 | pMBA$_{fosE}$ | *fosE* (mix) | “ |
| A624 | MG1655 | pMBA$_{fosF}$ | *fosF* (mix) | “ |
| A354 | MG1655 | pMBA$_{fosG}$ | *fosG* (mix) | “ |
| A355 | MG1655 | pMBA$_{fosH}$ | *fosH* (mix) | “ |
| A356 | MG1655 | pMBA$_{fosI}$ | *fosI* (mix) | “ |
| A626 | MG1655 | pMBA$_{fosK}$ | *fosK* (mix) | “ |
| A621 | MG1655 | pMBA$_{fosL}$ | *fosL* (mix) | “ |
| A622 | MG1655 | pMBA$_{fosM}$ | *fosM* (mix) | “ |
| A627 | MG1655 | pMBA$_{fosN}$ | *fosN* (mix) | “ |
| A337 | MG1655 | pMBA$_{smr1}$ | *smr1* (mix) | “ |
| A359 | MG1655 | pMBA$_{smr2}$ | *smr2* (mix) | “ |
| A366 | MG1655 | pMBA$_{smr3}$ | *smr3* (mix) | “ |
| A367 | MG1655 | pMBA$_{qacE}$ | *qacE* (mix) | “ |
| B091 | MG1655 | pMBA$_{qacE\Delta sul1}$ | *qacEΔsul1* (mix) | “ |
| C382 | DH5a | pMBA$_{qacE\Delta sul1}$ | *qacEΔsul1* (mix) | “ |
| A323 | MG1655 | pMBA$_{qacF}$ | *qacF* (mix) | “ |
| A357 | MG1655 | pMBA$_{qacG}$ | *qacG* (mix) | “ |
| A324 | MG1655 | pMBA$_{qacH}$ | *qacH* (mix) | “ |
| A623 | MG1655 | pMBA$_{qacK}$ | *qacK* (mix) | “ |
| A325 | MG1655 | pMBA$_{qacL}$ | *qacL* (mix) | “ |
| A358 | MG1655 | pMBA$_{qacM}$ | *qacM* (mix) | “ |
| **Quantitation of the cost of GFP expression** | | | | |
| A696 | MG1655 | pMBA ΔmetGFP KO | - | This study |
| A744 | MG1655 | pMBA$_{aacA59}$ ΔmetGFP KO | *aacA59* (aa) | “ |
| A745 | MG1655 | pMBA$_{aacA8}$ ΔmetGFP KO | *aacA8* (aa) | “ |
| A746 | MG1655 | pMBA$_{aadB}$ ΔmetGFP KO | *aadB* (aa) | “ |
| A747 | MG1655 | pMBA$_{aacC2}$ ΔmetGFP KO | *aacC2* (aa) | “ |
| A748 | MG1655 | pMBA$_{aacA28}$ ΔmetGFP KO | *aacA28* (aa) | “ |
| A749 | MG1655 | pMBA$_{aacA52}$ ΔmetGFP KO | *aacA52* (aa) | “ |
| A750 | MG1655 | pMBA$_{aacA30}$ ΔmetGFP KO | *aacA30* (aa) | “ |
| A751 | MG1655 | pMBA$_{aadA24}$ ΔmetGFP KO | *aadA24* (aa) | “ |
| A752 | MG1655 | pMBA$_{aadA13}$ ΔmetGFP KO | *aadA13* (aa) | “ |
| A753 | MG1655 | pMBA$_{aacC13}$ ΔmetGFP KO | *aacC13* (aa) | “ |
| A754 | MG1655 | pMBA$_{dfrA35}$ ΔmetGFP KO | *dfrA35* (dfr) | “ |
| A755 | MG1655 | pMBA$_{dfrB4}$ ΔmetGFP KO | *dfrB4* (dfr) | “ |
| A756 | MG1655 | pMBA$_{dfrA22.2}$ ΔmetGFP KO | *dfrA22.2* (dfr) | “ |
| A757 | MG1655 | pMBA$_{dfrA16}$ ΔmetGFP KO | *dfrA16* (dfr) | “ |
| A758 | MG1655 | pMBA$_{dfrA14}$ ΔmetGFP KO | *dfrA14* (dfr) | “ |
| A759 | MG1655 | pMBA$_{dfrA34}$ ΔmetGFP KO | *dfrA34* (dfr) | “ |
| **Mice and long-term competitions** | | | | |
| B878 | MG1655 | pMBA ΔGFP | - | “ |
| B880 | MG1655 | pMBA$_{ereA2}$ ΔGFP | *ereA2* (mix) | “ |
| B881 | MG1655 | pMBA$_{dfrA31}$ ΔGFP | *dfrA31* (dfr) | “ |
| B882 | MG1655 | pMBA$_{dfrA21}$ ΔGFP | *dfrA21* (dfr) | “ |
| B883 | MG1655 | pMBA$_{aacA7}$ ΔGFP | *aacA7* (aa) | “ |
| B886 | MG1655 | pMBA$_{bla\mathrm{OXA\text{-}10}}$ ΔGFP | *bla*$_{\mathrm{OXA\text{-}10}}$ (bla) | “ |
| 26074 | YL2 | - | - | *Brugiroux et al,* 2016 |
| 28989 | YL27 | - | - | “ |
| 26085 | I48 | - | - | “ |
| 26109 | YL45 | - | - | “ |
| 26127 | YL44 | - | - | “ |
| 32036 | KB1 | - | - | “ |
| 32035 | I49 | - | - | “ |
| 26114 | YL32 | - | - | “ |
| 26115 | YL58 | - | - | “ |
| 26117 | YL31 | - | - | “ |
| 26090 | KB18 | - | - | “ |
| 26113 | I46 | - | - | “ |

**Supplementary Table 8. Primers and probes used in this study.**

| Primer/Probe | Sequence (5´→ 3´) | Description |
|---|---|---|
| *int* R bb | CTTTGTTTTAGGGCGACTGC | pMBA linearisation |
| *gfp* F bb | TTAGGCGTCGACGCTGCA | |
| gBlock F | GCAGTCGCCCTAAAACAAAG | ARC amplification |
| gBlock R | GCTGCAGCGTCGACGCCTAA | |
| *int* F | AACGCAATTACAGAAATGCCTCGACTTCGC | pMBA sequencing |
| *gfp* R | AAAGCGCTGTCTAGACTATT | |
| *gfp* 2.0 R | CTCGATTCTATTAACAAGGG | |
| Δmet *gfp* F | GTAAATATAGAGTAAATGAGAAGAACTTTT | pMBA gfp KO and Δmet |
| Δmet *gfp R* | ATTTACTCTATATTTACCTCCTTTATATTG | |
| Δ*gfp F* | TGCAATATAAAATTGTCGGCACGTAAGAGG | pMBA *gfp* deletion |
| Δ*gfp R* | GCCGACAATTTTATATTGCAATCGCTGCAG | |
| Isol46 Exonucl.2 F | CGGATCGTAAAGCTCTGTTGTAAG | I46 qPCR (*Brugiroux et al*, 2016) |
| Isol46 Exonucl.3 R | GCTACCGTCACTCCCATAGCA | |
| Probe3 Isol46 | FAM-AAGAACGGCTCATAGAGG-BHQ1 | |
| Isol49 Exonucl. F | GCACTGGCTCAACAGATTGATG | I49 qPCR (*Brugiroux et al*, 2016) |
| Isol49 Exonucl. R | CCGCCACTCACTGGTGATC | |
| Probe Isol49 | HEX-CTTGCACCTGATTGACGA-BHQ1 | |
| YL58 Exonucl. F | GAAGAGCAAGTCTGATGTGAAAGG | YL58 qPCR (*Brugiroux et al*, 2016) |
| YL58 Exonucl. R | CGGCACTCTAGAAAAACAGTTTCC | |
| Probe YL58 | FAM-TAACCCCAGGACTGCAT-BHQ1 | |
| YL27 Exonucl.2 F | TCAAGTCAGCGGTAAAAATTCG | YL27 qPCR (*Brugiroux et al*, 2016) |
| YL27 Exonucl.2 R | CCCACTCAAGAACATCAGTTTCAA | |
| Probe2 YL27 | HEX-CAACCCCGTCGTGCC-BHQ1 | |
| YL31 Exonucl.2 F | AGGCGGGATTGCAAGTCA | YL31 qPCR (*Brugiroux et al*, 2016) |
| YL31 Exonucl.3 R | CCAGCACTCAAGAACTACAGTTTCA | |
| Probe2 YL31 | FAM-CAACCTCCAGCCTGC-BHQ1 | |
| YL32 Exonucl.2 F | AATACCGCATAAGCGCACAGT | YL32 qPCR (*Brugiroux et al*, 2016) |
| YL32 Exonucl.2 R | CCATCTCACACCACCAAAGTTTT | |
| Probe2 YL32 | HEX-CGCATGGCAGTGTGT-BHQ1 | |
| KB1 Exonucl. F | CTTCTTTCCTCCCGAGTGCTT | KB1 qPCR (*Brugiroux et al*, 2016) |
| KB1 Exonucl. R | CCCCTCTGATGGGTAGGTTACC | |
| Probe KB1 | FAM-CACTCAATTGGAAAGAGGAG-BHQ1 | |
| YL2 Exonucl. F | GGGTGAGTAATGCGTGACCAA | YL2 qPCR (*Brugiroux et al*, 2016) |
| YL2 Exonucl. R | CGGAGCATCCGGTATTACCA | |
| Probe2 YL2 | HEX-CGGAATAGCTCCTGGAAA-BHQ1 | |
| KB18 Exonucl.2 F | TGGCAAGTCAGTAGTGAAATCCA | KB18 qPCR (*Brugiroux et al*, 2016) |
| KB18 Exonucl.2 R | TCACTCAAGCTCGACAGTTTCAA | |
| Probe2 KB18 | FAM-CTTAACCCATGAACTGC-BHQ1 | |
| YL44 Exonucl. F | CGGGATAGCCCTGGGAAA | YL44 qPCR (*Brugiroux et al*, 2016) |
| YL44 Exonucl. R | GCGCATTGCTGCTTTAATCTTT | |
| Probe YL44 | HEX-TGGGATTAATACCGCATAGTA-BHQ1 | |
| YL45 Exonucl. F | AGACGGCCTTCGGGTTGTA | YL45 qPCR (*Brugiroux et al*, 2016) |
| YL45 Exonucl. R | CGTCATCGTCTATCGGTATTATCAA | |
| Probe YL44 | FAM-ACCACTTTTGTAGAGAACGA-BHQ1 | |
| Isol48 Exonucl. F | GGCAGCATGGGAGTTTGCT | I48 qPCR (*Brugiroux et al*, 2016) |
| Isol48 Exonucl. R | TTATCGGCAGGTTGGATACGT | |
| Probe Isol48 | HEX-CAAACTTCCGATGGCGAC-BHQ1 | |
| *E. coli* Exonucl. F | GGACCTTCGGGCCTCTTG | *E. coli* qPCR (*Brugiroux et al*, 2016) |
| *E. coli* Exonucl. R | CCTTTACCCCACCTACTAGCTAATCC | |
| Probe *E. coli* | FAM-ATCGGATGTGCCCAGAT-BHQ1 | |
