## Supplementary material for "Fitness effects of antimicrobial resistance genes in changing environments": Mathematical model

### Computational model of ARC stability

We developed a stochastic resource-explicit model to describe the competitive dynamics between two *E. coli* populations in a well-mixed, resource-limited environment. The reference population ($B_0$) carries the empty plasmid `pMBA∅`, whereas the competing population ($B_x$) carries a plasmid-borne antibiotic resistance cassette (ARC). The model captures how these populations grow and compete for a shared limiting resource under static or fluctuating oxygen regimes (aerobiosis $\leftrightarrow$ anaerobiosis).

The system state is represented by the vector

$$\mathbf{X}(t) = [B_0(t), B_x(t), R(t)],$$

where $B_0(t)$ and $B_x(t)$ denote the population sizes of the control and ARC-bearing strains, respectively, and $R(t)$ is the concentration of a single limiting resource. Growth of each strain follows Monod kinetics, such that the instantaneous division rate of strain $i \in \{0, x\}$ in environment $e$ is given by

$$r_i(e,R) = V_i(e)\,\frac{R}{R+K_i(e)},$$

where $V_i(e)$ is the maximum division rate and $K_i(e)$ the half-saturation constant. Each division consumes $c_i(e)$ units of the resource. The parameters $\{V_i(e), K_i(e), c_i(e)\}$ depend on the environmental condition, which can alternate between aerobic (E) and anaerobic (G) states.

The model assumes a homogeneous, well-mixed environment with no spatial structure or mutations. Population sizes are treated as discrete variables, and extinction is applied deterministically at the end of each simulated day: if $B_i < 1$, the lineage is considered extinct and its abundance is set to zero before dilution.

Cell division events are represented as Poisson processes whose expected number of occurrences within each time step $\Delta t$ is proportional to the instantaneous growth propensity. The model is implemented using a discrete $\tau$-leaping (Gillespie-like) scheme, in which multiple events may occur within each leap while propensities are assumed constant.

**Table 1.** Reaction channels, propensities, and state-change vectors. The environmental regime $e \in \{\mathrm{E}, \mathrm{G}\}$ modulates the parameters $\{V_i(e), K_i(e), c_i(e)\}$. The system state is $\mathbf{X}(t) = [B_0, B_x, R]^\top$.

| Index | Reaction | Propensity $a_j(\mathbf{X})$ | Change $\nu_j$ |
|---|---|---|---|
| $j=0$ | $B_0 \to B_0+1,\ R \to R-c_0(e)$ | $a_0 = V_0(e)\,B_0\,\frac{R}{R+K_0(e)}$ | $\nu_0 = [+1,\ 0,\ -c_0(e)]^\top$ |
| $j=1$ | $B_x \to B_x+1,\ R \to R-c_x(e)$ | $a_1 = V_x(e)\,B_x\,\frac{R}{R+K_x(e)}$ | $\nu_1 = [0,\ +1,\ -c_x(e)]^\top$ |

At each time step $\Delta t$, independent Poisson deviates are drawn for each reaction channel:

$$n_j \sim \mathrm{Poisson}\big(a_j(\mathbf{X}(t))\,\Delta t\big), \qquad j \in \{0,1\},$$

and the state vector is updated according to

$$\mathbf{X}(t+\Delta t) = \mathbf{X}(t) + \sum_{j=0}^{1} n_j\,\nu_j.$$

Expanding these updates explicitly for each strain yields

$$\lambda_{\mathrm{birth},i} = V_i(e)\,B_i(t)\,\frac{R(t)}{R(t)+K_i(e)}\,\Delta t, \tag{1}$$

$$n_{\mathrm{birth},i} \sim \mathrm{Poisson}\big(\lambda_{\mathrm{birth},i}\big), \tag{2}$$

$$B_i(t+\Delta t) = B_i(t) + n_{\mathrm{birth},i}, \tag{3}$$

$$R(t+\Delta t) = R(t) - \sum_i c_i(e)\,n_{\mathrm{birth},i}. \tag{4}$$

All variables are constrained to remain non-negative after each update:

$$B_i(t) \geq 0, \qquad R(t) \geq 0.$$

If a population falls below one cell at the end of a daily cycle, it is considered extinct and removed from subsequent updates.

In the limit of large populations, where stochastic fluctuations become negligible, the model reduces to the deterministic Monod system:

$$\frac{dB_i}{dt} = V_i(e)\, B_i \,\frac{R}{R + K_i(e)}, \tag{5}$$

$$\frac{dR}{dt} = -\sum_i c_i(e)\, V_i(e)\, B_i \,\frac{R}{R + K_i(e)}. \tag{6}$$

This deterministic approximation serves as a reference for the expected dynamics of the stochastic formulation when demographic noise is small.

### Model parameterization

Model parameters were derived directly from experimental measurements and used to construct a large synthetic library of antibiotic resistance cassettes (ARCs) with realistic fitness effects. Parameterization proceeded in three main steps: calibration of the reference strain, estimation of family-level fitness distributions from experiments, and generation of synthetic competitors consistent with these distributions.

The reference strain ($B_0$), carrying the empty plasmid $\texttt{pMBA}_\emptyset$, was first characterized in monoculture in aerobiosis. Its parameters were estimated by fitting the deterministic form of the model to experimental growth curves. The parameter vector

$$p_0 = \{V_0, K_0, c_0\}$$

defines the maximum division rate ($V_0$), the half-saturation constant of the limiting resource ($K_0$), and the amount of resource consumed per cell division ($c_0$). The fitted values accurately reproduced the empirical growth trajectories and served as the baseline for all subsequent simulations.

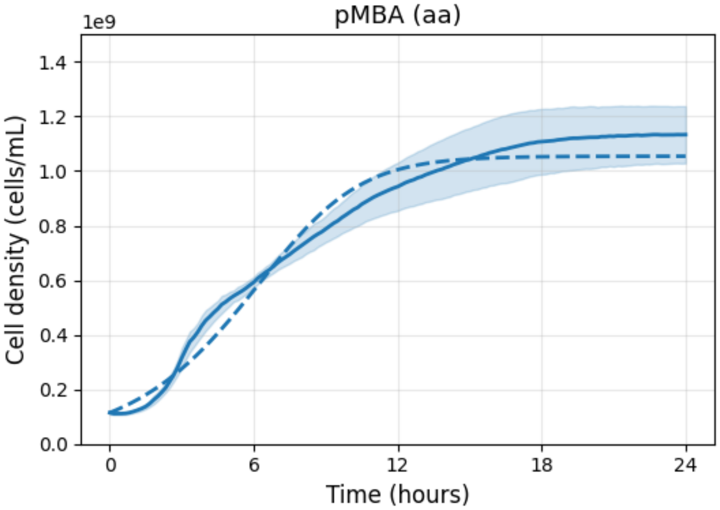


**Figure 1. Parameterization of the reference strain.** Calibration of the growth parameters for the reference strain carrying the empty plasmid ($\texttt{pMBA}_\emptyset$) in monoculture. Experimental growth curves (solid line) were obtained from replicate optical density measurements and represent mean values with associated standard deviations. The simulated trajectory (dotted line) corresponds to the best fit of the deterministic model using the parameters $\{V_0, K_0, c_0\}$.

Experimental competitions between ARC-bearing and control strains provided direct estimates of the relative fitness $w$ for each cassette under aerobic and anaerobic conditions. For each ARC family (`aa`, `bla`, `dfr`, `mix`), these paired measurements defined a joint empirical distribution in the two-dimensional fitness space $(w^{\mathrm{E}}, w^{\mathrm{G}})$. To capture both the variance and the observed correlation between environments, the family-level distributions were modeled as bivariate normal:

$$\begin{bmatrix} w^{\mathrm{E}} \\ w^{\mathrm{G}} \end{bmatrix} \sim \mathcal{N}_2\left( \begin{bmatrix} \mu_{\mathrm{E}} \\ \mu_{\mathrm{G}} \end{bmatrix}, \begin{bmatrix} \sigma_{\mathrm{E}}^2 & \rho\, \sigma_{\mathrm{E}}\sigma_{\mathrm{G}} \\ \rho\, \sigma_{\mathrm{E}}\sigma_{\mathrm{G}} & \sigma_{\mathrm{G}}^2 \end{bmatrix} \right),$$

where $\mu_{\mathrm{E}}$ and $\mu_{\mathrm{G}}$ are the family-specific mean fitnesses in aerobic and anaerobic environments, $\sigma_{\mathrm{E}}$ and $\sigma_{\mathrm{G}}$ are the corresponding standard deviations, and $\rho$ is the empirical correlation coefficient between them.

For each simulated ARC, a pair of target fitness values $(w_*^{\mathrm{E}}, w_*^{\mathrm{G}})$ was drawn jointly from the bivariate distribution. A parameter vector

$$p_x = \{V_x, K_x, c_x\}$$

was then determined separately for each environment (E and G) such that the simulated fitness

$$w_{\mathrm{sim}}(p_0, p_x)$$

approximated the sampled target value in that condition. The simulated fitness was defined as the ratio of growth parameters between the ARC-bearing and reference strains in short-term numerical competitions.

By repeating this calibration, we generated a synthetic population of 1000 ARC-bearing competitors per family. Experimental ARCs were included explicitly, and additional strains were drawn to complete the joint distribution (Supplementary Figure 2). This approach ensures that simulated fitness values reproduce not only the empirical mean and variance of each family but also the covariance between environmental responses, yielding family-specific fitness landscapes we will use for the subsequent numerical simulations.

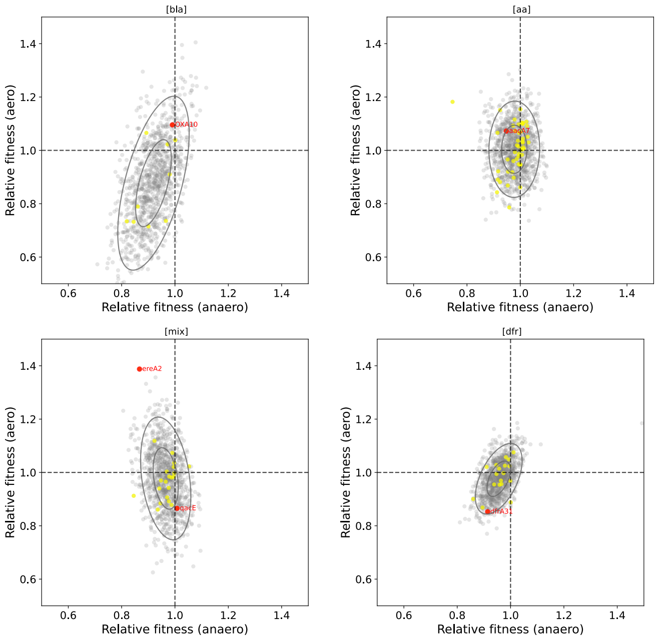


**Figure 2. Bivariate distributions of ARC fitness across environments.** Joint and marginal distributions of relative fitness ($w$) measured under anaerobic (x-axis) and aerobic (y-axis) conditions for all ARC families. Each point represents an individual ARC, with shaded ellipses indicating the 95% confidence contours of the bivariate normal fits for each family. Dashed lines mark neutrality ($w = 1$). (A) `bla` family, (B) `aa` family, (C) `mix` family, and (D) `dfr` family.

### Numerical experiments

Each simulation consisted of a sequence of 24-hour growth cycles followed by dilution, mimicking a serial-transfer experiment. During each cycle, the system evolved according to the stochastic update rules described above, with integration proceeding in fixed time steps of $\Delta t = 0.1\,\text{h}$ until 24 h of simulated time were completed. At the end of each simulated day, extinction was evaluated: if $B_i < 1$, the lineage was considered extinct and its abundance was set to zero. Surviving populations were then diluted by a constant factor $d$ (typically 1:100) and the resource concentration was reset to its initial value $R_0$. This growth–dilution process was repeated for $n_{\text{transfers}} = 100$ consecutive days.

The environmental state $e(t)$ determined which parameter set $\{V_i(e), K_i(e), c_i(e)\}$ applied during a given day. Three general regimes were considered: constant aerobiosis ($e = \text{E}$), constant anaerobiosis ($e = \text{G}$), and alternating oxygen levels following a prescribed switching schedule $e(t)$. Environmental schedules were implemented as discrete daily sequences defined by a single integer parameter $K$, representing the number of consecutive anaerobic days before the environment switched to aerobic conditions. In this scheme, $K = 0$ corresponds to constant anaerobiosis, $K = n_{\text{days}}$ to constant aerobiosis, and intermediate values $0 < K < n_{\text{days}}$ describe single-switch regimes from $\text{G} \to \text{E}$. Formally, the environmental state can be expressed as

$$e(t) = \begin{cases} \text{G}, & t \le K, \\ \text{E}, & t > K. \end{cases}$$

Simulations were initialized with both populations at equal density,

$$B_0(0) = B_x(0) = B_{\text{init}},$$

and with the resource concentration set to

$$R(0) = R_0.$$

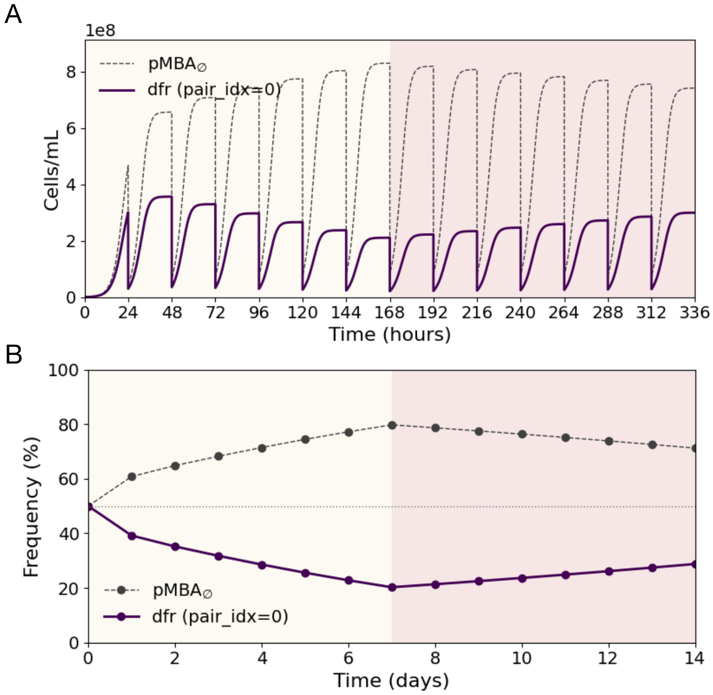


**Figure 3. Stochastic growth–dilution simulation of pairwise competition.** (A) Time-resolved population trajectories of the reference strain (`pMBA∅`) and a representative ARC-bearing strain over multiple 24 h growth cycles. Cell density (cells/mL) is shown on the $y$ axis, and time (hours) on the $x$ axis. The colored band indicates the environmental state applied on each day (yellow: anaerobic, light-red: aerobic). (B) Daily endpoints of the same simulation, expressed as the relative frequency of the ARC-bearing strain at the end of each transfer. The same environmental schedule is shown as a background band.

Each competition (ARC pair × environment × schedule) was simulated independently for several stochastic replicates ($n = 3$ by default). Daily trajectories were then summarized as replicate averages,

$$\langle B_i(t) \rangle = \frac{1}{n} \sum_{r=1}^{n} B_i^{(r)}(t),$$

which were used to reduce stochastic noise in long-term simulations.

At the end of each experiment, outcomes were classified according to the final frequency of the ARC-bearing strain,

$$f_{\mathrm{end}} = \frac{B_x}{B_x + B_0}.$$

Populations were categorized as cleared ($f_{\mathrm{end}} < 0.05$), fixated ($f_{\mathrm{end}} > 0.95$), or in stable coexistence ($0.05 \leq f_{\mathrm{end}} \leq 0.95$). Persistence was defined as the condition $f_{\mathrm{end}} > 0.01$, and a binary flag $p_{\mathrm{end}}$ was stored for each simulation:

$$p_{\mathrm{end}} = \begin{cases} 1, & \text{if } f_{\mathrm{end}} > 0.05, \\ 0, & \text{otherwise.} \end{cases}$$

The total simulated time was $T = 24\,\mathrm{h} \times n_{\mathrm{transfers}}$. For each ARC and environmental schedule, the simulations generated time series of $B_0(t)$, $B_x(t)$, and $R(t)$, together with daily summaries, final abundances, classification labels, and metadata identifying the ARC family, environmental regime, and replicate ID.

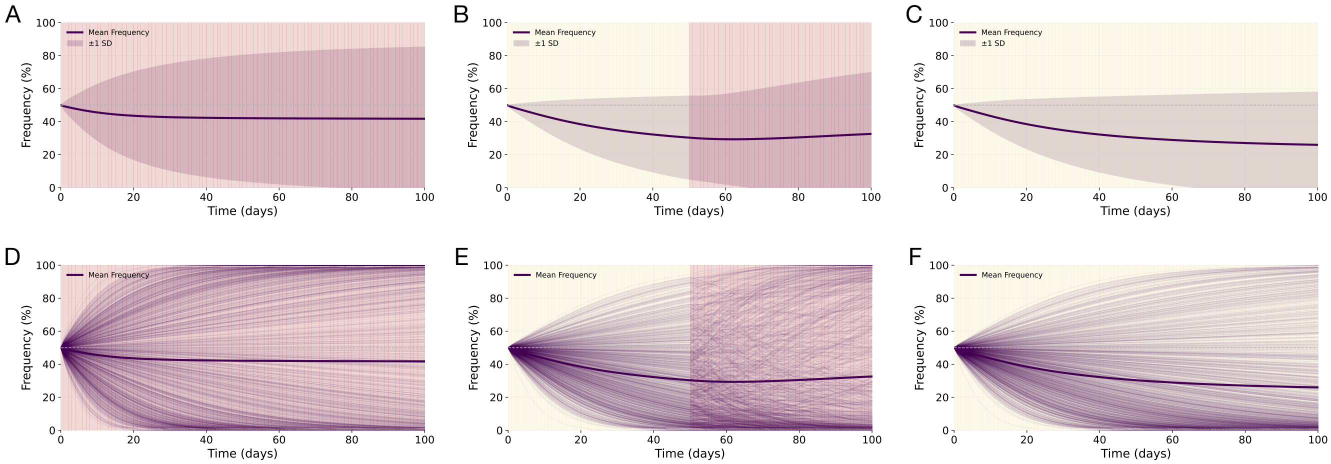


**Figure 4. ARC population dynamics under constant and switching environments.** Time-resolved frequencies of ARC-bearing strains relative to the reference strain ($\texttt{pMBA}_{\emptyset}$) over a 100-day serial-transfer simulation. Each column corresponds to a distinct environmental regime: constant anaerobiosis (A, D), a single switch from anaerobiosis to aerobiosis after $K = 40$ days (B, E), and constant aerobiosis (C,F). The upper panels (A-C) show the mean relative frequency of ARCs across the synthetic population ($n = 1000$) with associated standard deviations. The lower panels (D-F) display the same averages (thick line) together with individual trajectories of representative ARCs drawn from the synthetic library (thin lines). Background shading indicates the environmental state: anaerobic (yellow) and aerobic (light red).

#### Quantitative metrics

The primary observable was the probability that an ARC-bearing lineage remained detectable after a given number of transfer cycles. For a switching duration $K$, the persistence probability was defined as

$$P_{\mathrm{end}}(K) = \frac{\#\{\text{ARCs with } f_{\mathrm{end}}(K) > 0.01\}}{\#\{\text{ARCs tested}\}},$$

where $f_{\mathrm{end}}(K)$ is the final frequency of each ARC under the environmental schedule characterized by parameter $K$. This probability was estimated both for the full pool of ARCs and separately for each family (`aa`, `bla`, `dfr`, `mix`).

In addition to the binary persistence indicator $p_{\mathrm{end}}$, each simulation was assigned one of three categorical outcomes: cleared ($f_{\mathrm{end}} < 0.05$), stable coexistence ($0.05 \le f_{\mathrm{end}} \le 0.95$), or fixated ($f_{\mathrm{end}} > 0.95$). For each ARC family and environmental condition, the relative frequency of these outcomes defined the composition of the simulated community at the end of the experiment.

To quantify the impact of environmental switching on ARC survival, we defined a rescue probability measuring the fraction of lineages that would go extinct under constant anaerobiosis but persisted when the environment eventually became aerobic:

$$P_{\mathrm{rescue}}(K) = \frac{\#\{\text{ARCs with } p_{\mathrm{end}}(K) = 1 \text{ and } p_{\mathrm{end}}(0) = 0\}}{\#\{\text{ARCs tested}\}}.$$

This metric was computed for each family and for the pooled population of ARCs, providing a quantitative measure of how fluctuating oxygen levels promote the recovery of otherwise non-persistent lineages.

#### Numerical implementation

All simulations were implemented in `Python` using standard scientific libraries (`NumPy`, `pandas`, `matplotlib`). The stochastic resource-explicit model was coded as a fixed–time step (*tau-leaping*) simulation with $\Delta t = 0.1\,\mathrm{h}$. Each competition between the reference and ARC-bearing strain was simulated independently for several stochastic replicates. All code and data necessary to reproduce these simulations are available in the GitHub repository: `https://github.com/ccg-esb-lab`.

**Table 2.** Model parameters and simulation constants. All quantities are dimensionless unless otherwise noted.

| Symbol | Description | Typical value / source |
|---|---|---|
| $V_i(e)$ | Maximum division rate of strain $i$ in environment $e$ | Calibrated from data |
| $K_i(e)$ | Half-saturation constant of strain $i$ in environment $e$ | Calibrated from data |
| $c_i(e)$ | Resource consumed per cell division | Calibrated from data |
| $R_0$ | Initial resource concentration | 1 (arbitrary units) |
| $B_{\mathrm{init}}$ | Initial cell count per strain | $10^4$ |
| $d$ | Dilution factor per transfer | 100 |
| $n_{\mathrm{transfers}}$ | Number of serial transfers | 100 |
| $\Delta t$ | Integration time step | $0.1\,\mathrm{h}$ |
| $e(t)$ | Environmental state (E or G) | Defined by schedule |
| $K$ | Length of anaerobic phase before switch | Variable (0–100 days) |
| $f_{\mathrm{end}}$ | Final frequency of ARC-bearing strain | Computed from simulation |
| $p_{\mathrm{end}}$ | Binary persistence indicator | Computed from $f_{\mathrm{end}}$ |
| $P_{\mathrm{end}}(K)$ | Persistence probability at switch length $K$ | Derived metric |
| $P_{\mathrm{rescue}}(K)$ | Rescue probability relative to $K = 0$ | Derived metric |
